# Human Medial Ganglionic Eminence Organoids Robustly Generate Parvalbumin Interneurons and Fast-Spiking Neurons and Reveal Migratory Deficits in *SLC6A1* Deficient Interneurons

**DOI:** 10.1101/2025.07.01.662594

**Authors:** Maria C. Varela, Miranda P. Walker, Jeyoon Bok, Emmanuel L. Crespo, Tyler Thenstedt, Leah Goldstein, Andrew M. Tidball, Yukun Yuan, Lori L. Isom, Jianping Fu, Michael Uhler, Jack M. Parent

**Affiliations:** Department of Neurology, University of Michigan Medical School, Ann Arbor, MI, USA; Neuroscience Graduate Program, University of Michigan Medical School, Ann Arbor, MI, USA; Department of Mechanical Engineering, University of Michigan, Ann Arbor, MI, USA; Department of Pharmacology, University of Michigan Medical School, Ann Arbor, MI, USA; Department of Biomedical Engineering, University of Michigan, Ann Arbor, MI, USA; Department of Cell & Developmental Biology, University of Michigan Medical School, Ann Arbor, MI, USA; Department of Biological Chemistry, University of Michigan Medical School, Ann Arbor, MI, USA; Michigan Neuroscience Institute, University of Michigan Medical School, Ann Arbor, MI, USA; VA Ann Arbor Healthcare System, Ann Arbor, MI, USA

**Keywords:** GABAergic, brain organoid, pluripotent stem cells, neuronal migration, forebrain development, multielectrode array, basket cells, axoaxonic cells

## Abstract

The medial ganglionic eminence (MGE) gives rise to parvalbumin (PV)-and somatostatin (SST)-expressing cortical interneurons essential for regulating cortical excitability. Although PV interneurons are linked to various neurodevelopmental and neurodegenerative disorders, reliably generating them from human pluripotent stem cells (hPSCs) has been extremely challenging. We present a robust, reproducible protocol for generating single-rosette MGE organoids (MGEOs) from hPSCs.

Transcriptomic analyses reveal that MGEOs exhibit MGE regional identity and faithfully model the developing human fetal MGE. As MGEOs mature, they generate abundant PV-expressing cortical interneurons, including putative basket and axoaxonic cells, at a scale not previously achieved *in vitro*. When fused with human cortical organoids (hCOs), these interneurons rapidly migrate into the hCOs, integrate into excitatory networks, and contribute to complex electrophysiological patterns and the emergence of large numbers of fast-spiking neurons. Using this model, we uncover a previously unreported migration deficit of MGE interneurons in a disease model of *SLC6A1* developmental and epileptic encephalopathy, offering potential insights into the developmental contributions to epileptogenesis. MGEOs thus offer a powerful *in vitro* approach for probing human MGE-lineage cortical and subcortical GABAergic neuron development, modeling various neuropsychiatric disorders, and advancing cell-based therapies for neurodevelopmental and neurodegenerative disorders.

**Highlights:**

- Generation of subpallial organoids highly enriched for MGE lineages
- MGE organoids (MGEOs) robustly produce parvalbumin-expressing cortical interneurons
- Complex network activity and fast-spiking neurons are generated in assembloids
- Impaired interneuron migration in *SLC6A1* knockout and patient-derived MGEOs

**Graphical abstract:** 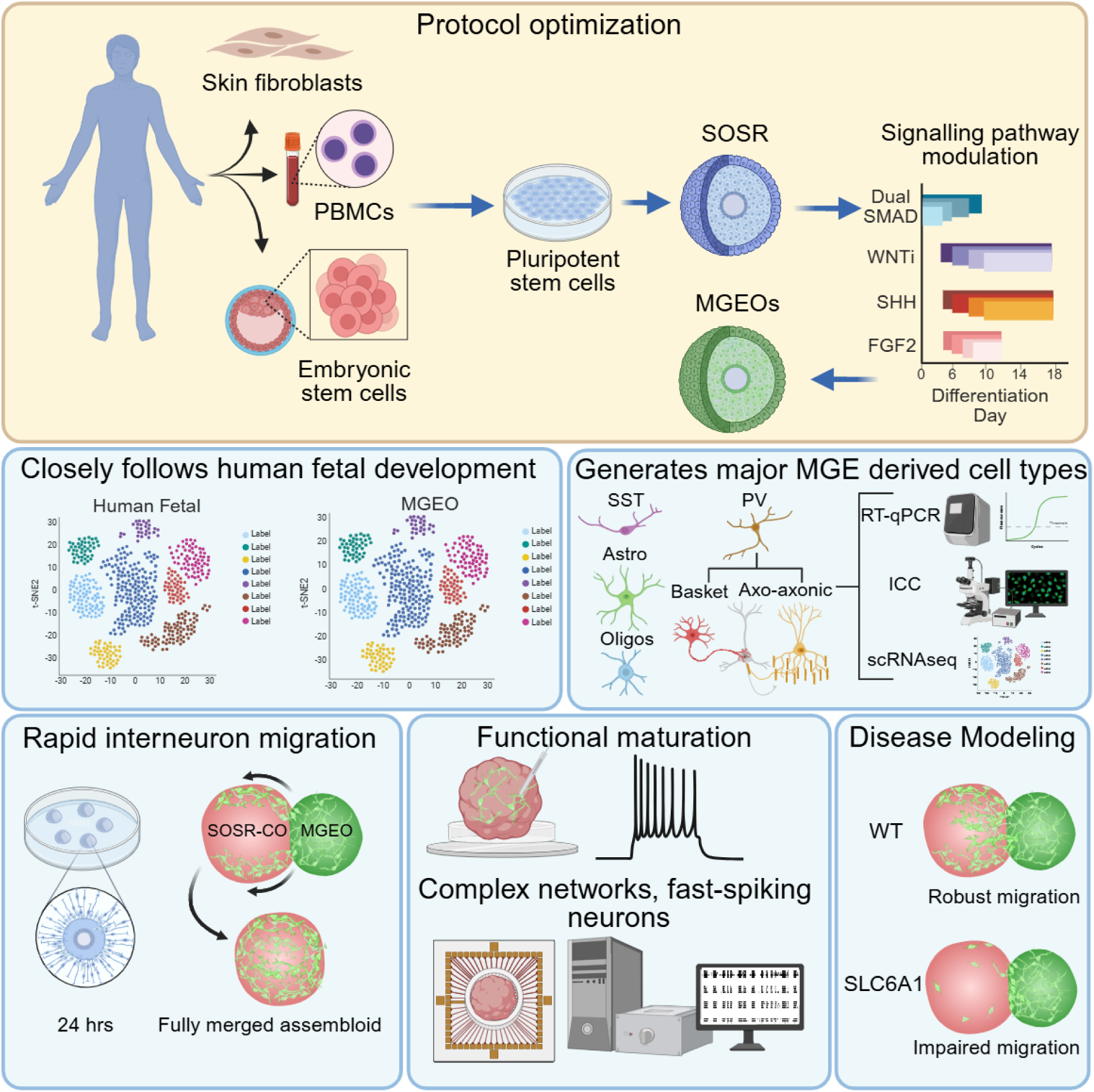

## INTRODUCTION

The human cortex is largely populated by two classes of neurons: glutamatergic excitatory neurons and gamma-aminobutyric acid (GABA)-containing inhibitory interneurons. During development, glutamatergic neurons are generated in the dorsal telencephalon and migrate radially as they form the cortical layers. In contrast, the majority of pallial GABAergic interneurons are generated in the ganglionic eminences (GEs) before migrating tangentially to the cortex. GABAergic neurons modulate brain network activity by regulating the timing and synchrony of neuronal firing, leading to the formation of neuronal oscillations critical for information transfer in the brain^1–4^.

Disruptions in the migration, maturation, synaptogenesis or function of these interneurons have been linked to various neurodevelopmental, neurologic and psychiatric disorders, such as autism, epilepsy, schizophrenia and Alzheimer’s disease ^5–19^, highlighting the critical nature of these cells in maintaining neural homeostasis.

The GEs are transient structures in the developing brain comprised of three distinct regions: medial ganglionic eminence (MGE), caudal ganglionic eminence (CGE) and lateral ganglionic eminence (LGE). The MGE and CGE, along with the preoptic area (POA), generate the majority of interneurons in the cortex and hippocampus. The MGE serves as the origin of approximately 60% of cortical interneurons and produces several interneuron subtypes crucial for cortical function^5–10^. The MGE is characterized by the expression of the forebrain marker FOXG1, as well as homeobox transcription factors NKX2.1 and LHX6^11–15^. *NKX2.1* is also expressed in the rostral pallidum, ventral septum, preoptic area, and ventral hypothalamic regions. In contrast, *LHX6* is largely restricted to MGE-derived lineages. In MGE-derived GABAergic lineages, upregulation of *LHX6* is induced by *NKX2.1* expression at approximately gestational week 8 (GW8) in humans or embryonic day 11.5 in mice and is essential for the proper positioning and maturation of cortical SST and PV interneurons ^5,16–24^. In addition to SST and PV cortical interneurons, the MGE also produces GABAergic and cholinergic striatal interneurons, subpallial GABAergic projection neurons, astrocytes, and oligodendrocytes^20,25,26^.

Recent transcriptomic studies of human fetal brain tissues have provided significant insights into the gene expression profiles of the developing human GEs^11,14,15,27,28^. However, the availability of human brain tissues from a broad range of fetal development is limited. To address this limitation, human pluripotent stem cell (hPSC)-derived 2D and 3D brain organoid models have emerged as promising tools for recapitulating neural development *in vitro*. In particular, brain organoids recapitulate key aspects of brain cytoarchitecture, cellular diversity, and complex network connectivity, and they offer the potential for unique insights into human neurodevelopment and associated disorders. Several pioneering groups have generated GABAergic neurons from hPSCs in both 2D cultures^12,13,29^ and brain organoid models^30–33^. These methods produce NKX2.1+ progenitors and generate primarily GABAergic SST+ neurons and neurons that arise from other subpallial regions but little to no PV+ cortical interneurons in culture.

PV+ interneurons are the largest class of interneurons in the human cortex, comprising approximately 30-40% of all cortical interneurons, and play a critical role in regulating cortical excitability. PV interneuron fast-spiking properties allow them to tightly regulate and synchronize cortical network activity through gamma oscillations. Dysregulation of PV interneuron development and function is implicated in epilepsy, schizophrenia, and autism spectrum disorders, making them a highly desirable cell type to generate for *in vitro* disease modeling^34–37^. However, ^51^the late generation and prolonged maturation time of PV interneurons has historically limited the field’s ability to derive and study them using hPSC models.

Recent studies have made progress in generating PV-expressing cortical basket interneurons and fast-spiking neurons from hPSCs using organoid assembloid^118^ approaches or xenotransplant strategies^46,119^. However, the efficiency in PV interneuron generation remains low, with PV interneuron clusters identified indirectly through predictive identity algorithms^119^, or via immunostaining that shows limited PV expression^118^. To date, no hPSC-derived *in vitro* model has shown robust, quantified expression of PV at the protein level. Furthermore, to our knowledge, PV-expressing axoaxonic cells have not been shown in any published human subpallial organoid or assembloid models.

Here we describe a human MGE-specific organoid (MGEO) model that exhibits MGE lineage marker expression and cell type specificity. The model exhibits transcriptomic similarities to human fetal MGE, generates highly migratory pallial interneuron progenitors, and displays robust development of MGE lineage interneuron subtypes that include SST and PV interneurons. When fused to human cortical organoids (hCOs) to generate assembloids, MGEO-derived interneurons migrate into hCOs where they integrate with excitatory cortical neurons, leading to complex network activity and the development of fast-spiking neurons. We also show the utility of MGEOs in disease modeling by identifying a novel, cell-autonomous MGE interneuron migration deficit in MGEO and assembloid models of *SLC6A1*-associated developmental and epileptic encephalopathy (DEE). Thus, our model offers a powerful platform for studying human-specific MGE interneuron development and disorders related to MGE interneuron dysfunction, and provides a potential source of human MGE interneuron progenitors for therapeutic grafting.

## RESULTS

### Timing and concentration of patterning molecules influence MGE specificity of subpallial organoids

Regional specification of the ganglionic eminences, including the MGE, is mediated by precisely controlled spatiotemporal signals of Wingless-related Integration Site (WNT), Sonic Hedgehog (SHH) and other patterning factors during forebrain development (**Figure 1A**). Established protocols for generating ventral telencephalic (subpallial) organoids rely on SHH pathway agonists and WNT pathway antagonists to guide organoids towards a ventral forebrain regional identity ^30,31,33^. We recently developed a method for generating dorsal forebrain patterned self-organizing single-rosette cortical organoids, termed SOSR-COs^38^. Given the robust cell-type specificity, reproducibility, and improved structural organization of single-rosette organoids for modeling early neurodevelopmental disorders,^38–40^ we used this approach to develop self-organizing single rosette MGEOs, with the aim of achieving MGE regional specificity with a high yield of PV+ cortical interneurons. To this end, we systematically altered the timing of dual SMAD inhibition, and the timing and concentrations of SAG and the WNT antagonist XAV939 (**Figure 1B**). As a readout, we quantified expression of on-and off-target region-specific mRNAs at 18 days in vitro (DIV) using RT-qPCR.

**Figure 1:**
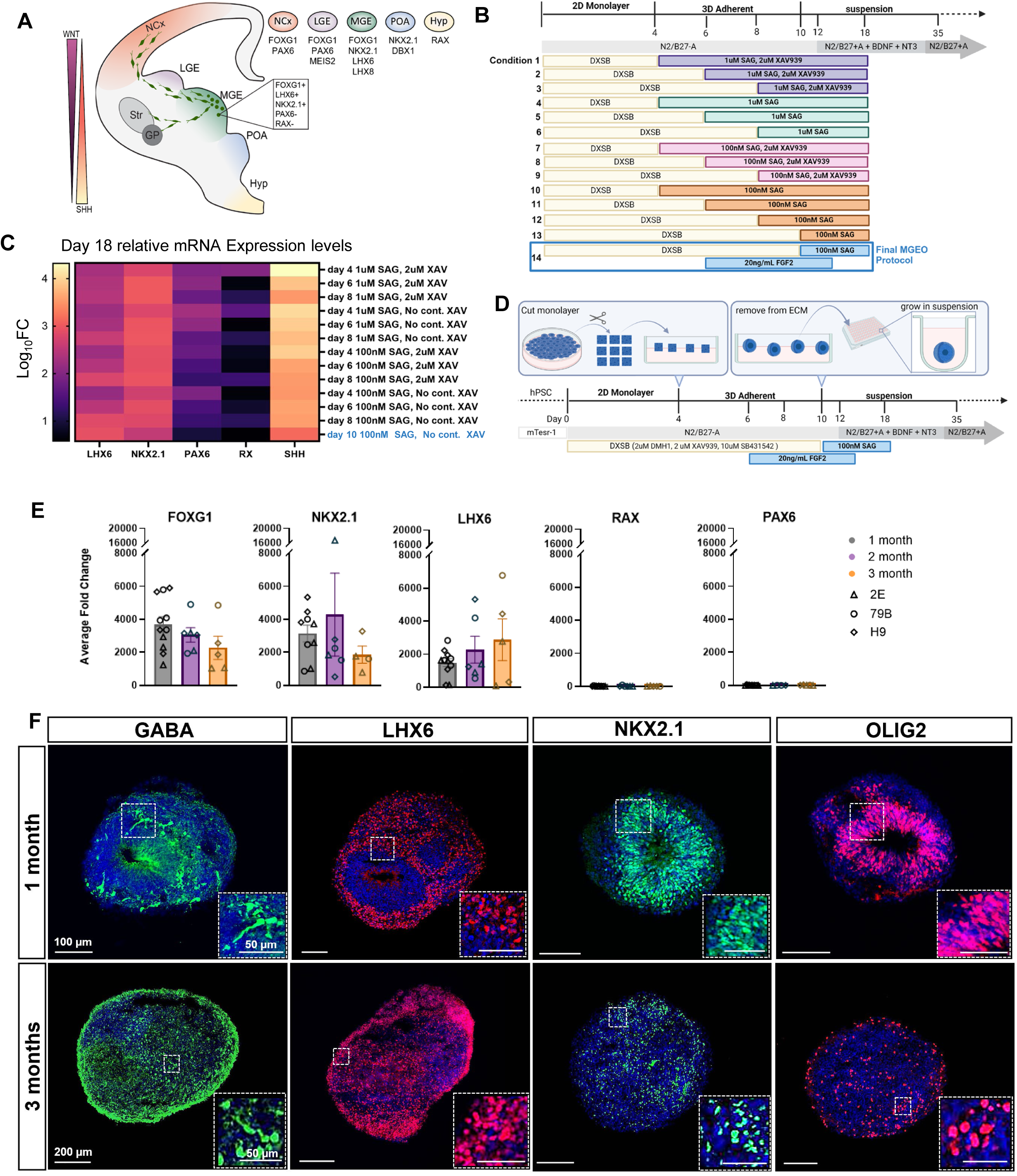
Generation of human subpallial brain organoids with MGE regional specificity (A) Schematic of MGE-derived interneuron tangential migration, key markers differentiating MGE from nearby regions, and the distribution of SHH and WNT patterning molecules. (B) Protocol variations tested for generating organoids with MGE specificity. DXSB condition: 2uM DMH1, 2uM XAV939, 10uM SB431542. (C) Heat map of relative mRNA levels measured by qRT-PCR in 18 DIV organoids exposed to various timings and concentrations of SAG with or without continued (cont.) XAV939 after DXSB. n=1-2 differentiations/condition of 2E iPSCs. Graphed as Log_10_FC relative to GAPDH and iPSC baseline. Blue text indicates the optimal conditions (high *LHX6*, low *PAX6* and *RX*). (D) Final MGEO protocol. Boxes in blue above the timeline illustrate the key steps at days 4 (monolayer cutting) and 10 (transition from 3D adherent to suspension culture). (E) Relative mRNA expression of 1-, 2-and 3-month MGEOs for key markers. n≥5 differentiations. Graphed as average ΔΔCT fold change ± SEM, normalized to GAPDH and relative to average iPSC baseline. Shapes denote hPSC line and colors indicate age (triangle: 2E, circle: 79B, diamond: H9; gray: 1 month, purple: 2 month, orange: 3 month). (F) MGEO immunostaining at 1-month (top) and 3-months (bottom) shows strong expression of GABA (green), LHX6 (red), NKX2-1 (green), and Olig2 (red). Bisbenzimide nuclear stain (DNA) in blue. Higher magnification views shown in insets. Abbreviations: NCx: Neocortex; LGE: Lateral Ganglionic Eminence; Str: Striatum; GP: Globus pallidus; MGE: medial ganglionic eminence; POA: preoptic area; Hyp: Hypothalamus; ECM: extracellular matrix.

We found that *PAX6*, a dorsal telencephalic marker with some expression in LGE and CGE, but not MGE^41^, was more highly expressed in conditions with earlier SAG exposure (**Figure 1C**). In contrast, *LHX6* expression was higher with later SAG exposure. Conditions with higher SAG concentrations resulted in greater levels of *SHH* mRNA expression, suggesting the generation of subpallial organoids with a more ventral telencephalic regional identity such as the POA or hypothalamus^42^. In contrast, *NKX2.1* was expressed at relatively similar levels across all conditions, consistent with its lack of MGE specificity. A combination of 100 nM SAG exposure from 10-18 DIV, brief XAV939 exposure during dual SMAD inhibition, and FGF2 treatment from 6-12 DIV showed the greatest MGE-like specificity as defined by high *LHX6* expression, moderate *NKX2.1* expression, and negligible levels of *PAX6* and the hypothalamic marker *RAX* (**Figure 1C**, bottom row in blue). We therefore selected this protocol for generating MGEOs for the remainder of the study (**Figure 1D**).

### MGEOs display high MGE regional specificity

To ensure the robustness of our protocol across different cell lines, we used human foreskin fibroblast-and blood-derived iPSC lines, denoted as 2E and 79B, respectively, and a human embryonic stem cell (hESC) line (H9/WA09). Continuing WNT inhibition until 10 DIV followed by SHH pathway activation from days 10 to 18 consistently produced MGEOs with robust expression of key forebrain-(*FOXG1*), ganglionic eminence progenitor-(*NKX2.1*), and MGE-specific (*LHX6*) markers for all the hPSC lines, as measured by RT-qPCR or immunocytochemistry from 1-3 months in vitro (**Figure 1E, F**)^43,44^. LHX6 mRNA and protein remained highly expressed in MGEOs from 1 to 3 months. Negligible mRNA expression levels of the hypothalamic marker RAX and the dorsal telencephalic marker *PAX6* were observed **(Figure 1E)**, with only miniscule amounts present in glial clusters of our scRNA-seq analyses **(Figures S2**), but no protein expression observed by immunocytochemistry for PAX6 or RAX in 1-month MGEOs (**Figure S1A-C**). Immunostaining of MGEOs at 1 and 3 months also showed the expected expression patterns for GABA and OLIG2 (**Figure 1F**).

To further validate the MGE specificity of our organoids, co-expression of OLIG2 and NKX2.1 was quantified by immunofluorescence analysis. In early developmental timepoints of the human brain (GW 8-12), OLIG2 is almost exclusively co-expressed with NKX2.1 in the MGE but not in surrounding brain regions including the LGE, CGE, hypothalamus, and POA^13,27^. At mid to late gestational weeks (GW 24+), NKX2.1 and OLIG2 segregate in their expression and do not co-express in mature MGE-derived cells. Our analysis of 1-month MGEOs shows a 98.5% colocalization of OLIG2+ cells with NKX2.1 (**Figure S1D, E**). In 3-month MGEOs, in contrast, most OLIG2+ cells did not co-express NKX2.1 (**Figure S1D, E**). These results further support the MGE specificity of our model.

We next performed single-cell RNA sequencing (scRNA-seq) analysis of 1-month MGEOs to assess their regional identity and cell type composition. Uniform Manifold Approximation and Projections (UMAP) plots revealed five distinct cell clusters annotated as radial glia (*VIM, NES, HES1*), inhibitory neuron progenitors (*ASCL1, HES6, DLX2*), and populations of GABAergic neurons that resemble tangentially migrating somatostatin (*SST*)+ cortical interneurons (*SST, LHX6, CXCR4, ERBB4, ZEB2, PLS3, ARX, MAF*), PV neuron precursors (*MAFB, MEF2C, ETV1, ST18, CRABP1*), and MGE subpallial destined neurons (*LHX8, GBX2*)^11,14,15,28^ (**Figure 2A-C**). Importantly, the percent distribution of each of these cell types in MGEOs was consistent within each sample replicate and between hPSC lines (**Figure 2B**). We confirmed the presence of cells in MGEOs expressing the MGE-specific marker LHX6, as well as the absence of cells expressing off-target markers that would indicate non-MGE identities, including CGE (*SP8, COUPTF2(NR2F2)*), LGE (*EBF1, ZFHX3*), hypothalamus (*NKX2-2, RAX*), POA (*HMX3, DBH*) and dorsal telencephalon (*PAX6*) (**Figures 2C and S2A**). Together, these data strongly support the early MGE specificity of MGEOs.

**Figure 2:**
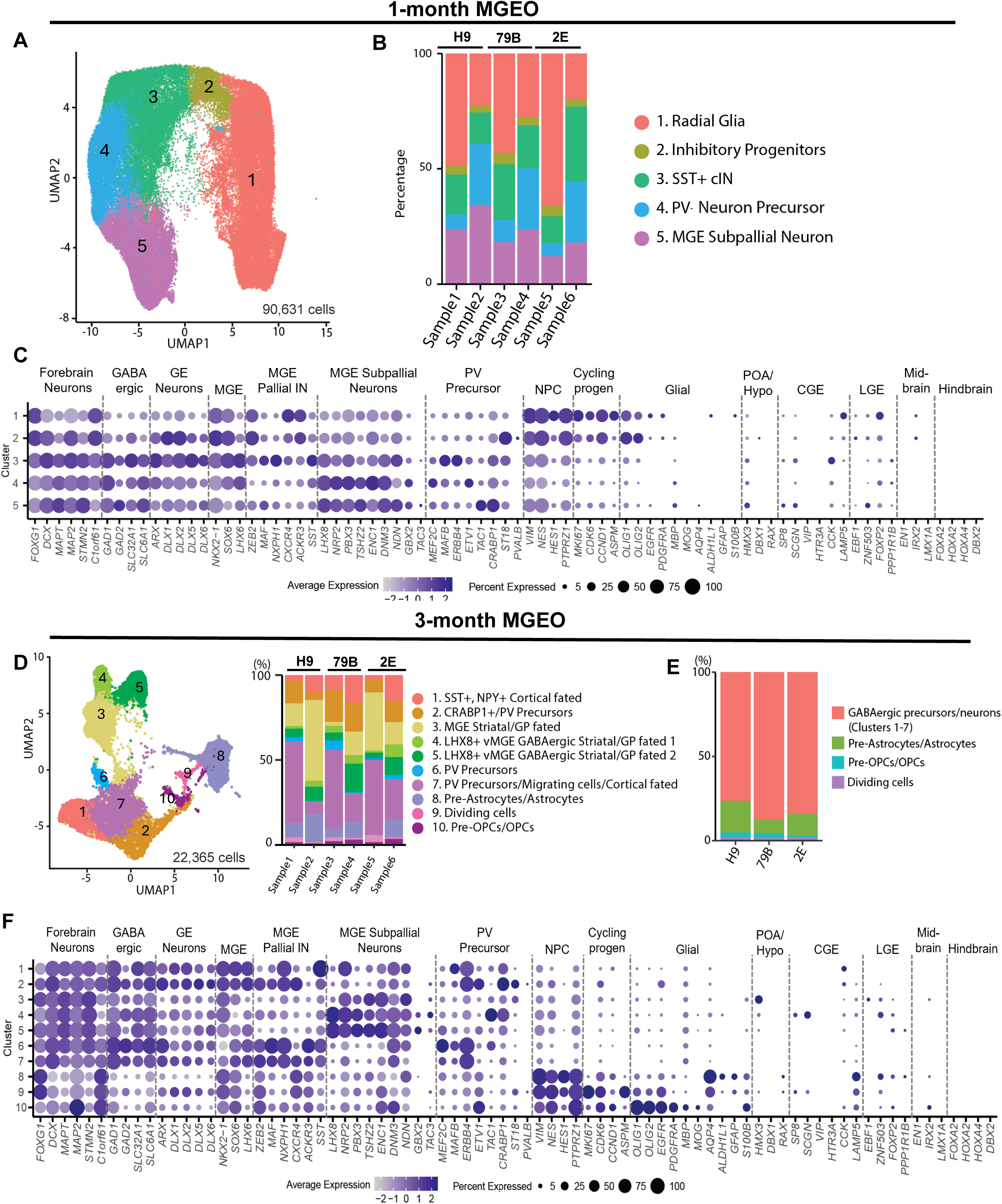
Single-Cell RNA sequencing of 1-month and 3-month MGEOs (A) UMAP plot of 1-month-old MGEOs with color coded and numbered clusters. n= 90,631 total cells from 6 samples (2 experimental replicates per hPSC line, 6-8 organoids pooled per sample). (B) Left: Stacked bar graphs denoting the cluster percent composition for each individual sample included in the UMAP in panel A. Right: Cluster name and color key. (C) Dot plot showing expression of markers of interest by cell cluster. Circle size denotes percent of cells within clusters expressing the gene of interest. Dot color indicates average expression level relative to other clusters. Dots were included for genes that contained > 1 percent of cells in a cluster. All cells with >0 expression levels were included. (D) UMAP plot shows 10 clusters from six 3-month-old MGEO samples, 2 differentiations/cell line, with clusters color-coded and numbered. Middle: bar graphs denote percentages of cells within each sample corresponding to clusters on the UMAP. Cells from 4-6 pooled organoids/sample from H9 (samples 1 & 2), 79B (samples 3 & 4), and 2E (samples 5 & 6) hPSC lines. Right: Cluster identities by number and color. (E) Average percentage of cells in each cluster by cell line. (F) Dot plots showing expression of markers of interest by 3-month MGEO cluster. Key and analysis as in C. Abbreviations: GE: Ganglionic Eminences; MGE: Medial Ganglionic Eminence; IN: Interneurons; POA: Preoptic area; CGE: Caudal Ganglionic Eminence; LGE: Lateral Ganglionic Eminence; NPC: Neural Progenitor Cell; Cycling Projen: Cycling Progenitors.

### MGEOs produce human MGE-derived cell types

The MGE primarily generates SST and PV GABAergic neuron subtypes, although there is evidence that to a lesser extent it generates neurons that express *CALB2* (calretinin), TH, and CHAT^5,43,45,46^. The MGE also gives rise to astrocytes and oligodendrocytes^47,48^. To further characterize the cell types produced in MGEOs, we conducted scRNAseq analysis at 3 months, RT-qPCR at 1-3 months, and immunostaining from 3-8 months in vitro. scRNA-seq data analysis of 3-month MGEOs generated from the three hPSC lines revealed distinct MGE-like cell lineages including: cortical PV precursors with or without CRAPBP1, SST+ cortical-destined cells, and MGE subpallial/striatal/globus pallidus-destined cells with or without *LHX8* expression (**Figure 2D**). We also identified astrocytes/pre-astrocytes and pre-oligodendrocyte precursor cells (pre-OPCs)/OPCs in the MGEOs (**Figure 2D, E**).

Although some variability was apparent in the cellular composition between experimental replicates and MGEOs generated from different hPSC lines (**Figure 2D**), the average percentages of GABAergic precursors/neurons, pre-astrocytes/astrocytes, pre-OPCs/OPCs, and dividing cells were highly consistent between cell lines and replicates (**Figure 2E**). As expected, there was negligible expression of the CGE lineage *VIP* or *CCK* GABAergic neuron subtypes in 3-month MGEOs (**Figure S3A, C**). Expression of *NPY* in 3-month MGEOs was limited to *SST*+/*NPY*+ cortical-destined neurons (**Figures 2D and S3A**). *CALB2*+ cells that co-expressed *NKX2.1* were found in the striatal/GP-destined clusters, and those that co-expressed *LHX6* appeared in cortical-destined clusters (data not shown). Neither of these populations substantially expressed the CGE markers *SP8* or *NR2F2* (*COUPTFII*) **(Figures 2C, S2B and S3B**), supporting non-CGE origins. Minimal expression of other off-target region-specific markers, such as POA/hypothalamus, midbrain and hindbrain, was observed (**Figure 2C, F**).

To confirm selected scRNA-seq findings, we next performed RT-qPCR for specific transcripts. Analysis of MGEOs revealed that the GABAergic neuronal markers VGAT and SST were expressed in MGEOs generated from all three hPSC lines at one, two, and three months (**Figure 3A**). Expression of *SLC12A5* (KCC2) relative to *SLC32A2* (NKCC1) progressively increased from one month to three months, consistent with neuronal maturation. As expected, *LHX8* expression was highly upregulated in MGEOs at 1 month and decreased thereafter. While calretinin (CR; *CALB2*)-expressing interneurons primarily originate from the CGE, a subpopulation is known to arise from the dorsal MGE^5,17,49,50^. We identified populations of CR-and SST-expressing neurons that co-expressed *LHX6*, indicating that both populations were derived from an MGE-like lineage (**Figure 3B, C**). Immunostaining of MGEOs at 3 months also revealed expression of GFAP in putative astrocytes, and sparse oligodendrocytes expressing both myelin basic protein (MBP) and OLIG2 (**Figure 3D**).

**Figure 3.**
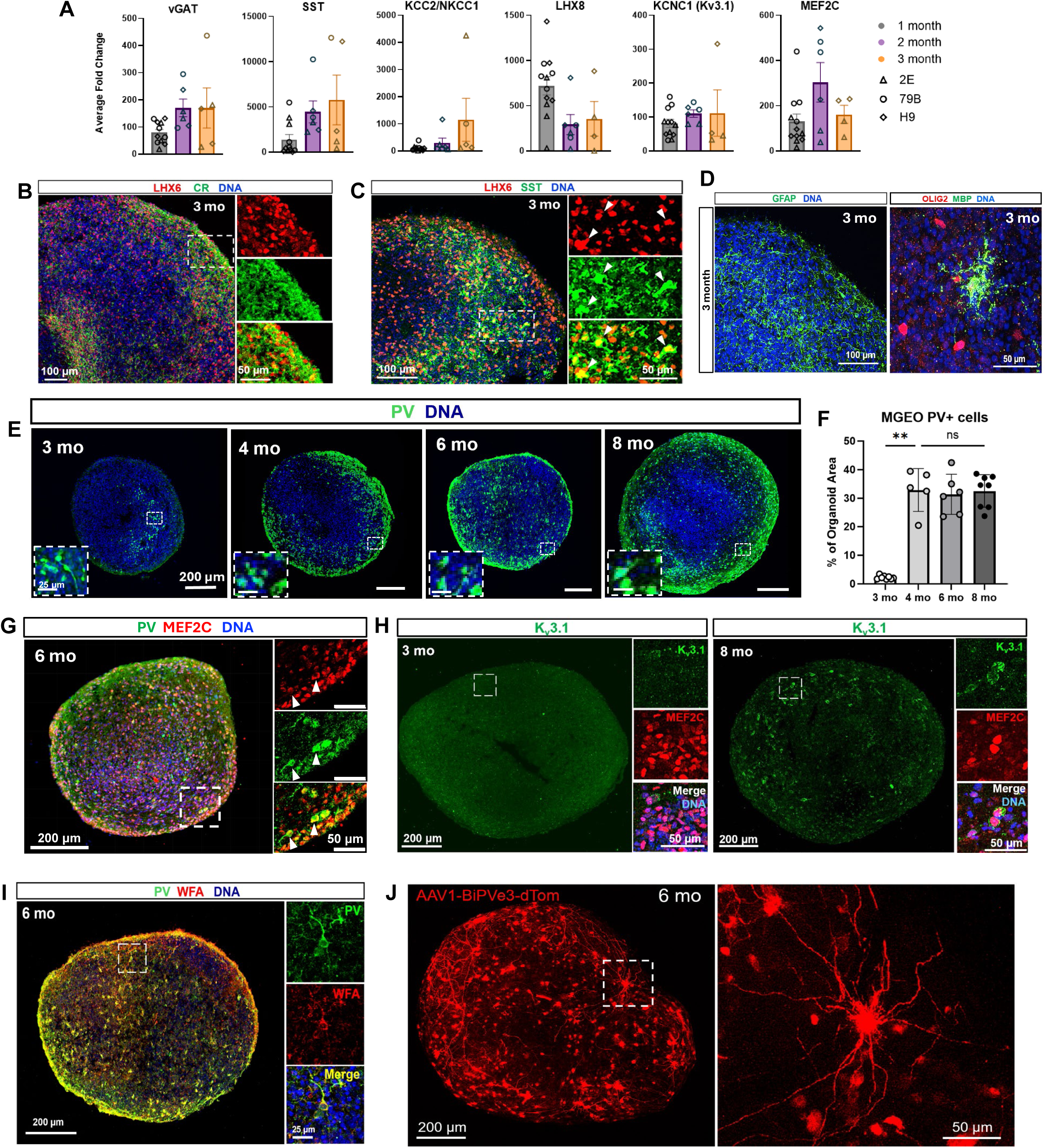
MGEOs generate human MGE-derived cell-types (A) RT-qPCR of 1-, 2-and 3-month MGEOs shows expression of GABAegic, chloride transporter, potassium channel, and MGE transcripts. n≥5 differentiations between 3 cell lines, graphed as average fold change (ΔΔCT) ± SEM, normalization to GAPDH and relative to average iPSC baseline. Shapes denote cell line and color indicates time point (triangle: 2E, circle: 79B, diamond: H9; gray: 1-month, purple: 2-month, orange: 3-month). (B-D) Representative confocal images of cell type-specific marker immunolabeling in 3-month (mo) MGEOs. Dashed boxes in B and C are shown in higher magnification on the right. DNA is bisbenzimide nuclear stain. (E) Representative confocal images of parvalbumin (PV, green) immunostaining at 3, 4, 6 and 8 months shows robust PV expression in MGEOs from 4 months onwards. Dashed boxes shown at higher magnification in lower left insets. (F) PV expression as percentage (%) of organoid area ± SD. n=5-8 organoids per timepoint and 2-3 differentiations/line. One way ANOVA with Dunnett’s T3 multiple comparisons post hoc test. **, P= 0.0036. ns, non-significant. (G) Immunostaining of 6-month organoids shows colocalization of PV (green) and MEF2C (red). Dashed box enlarged at right. (H) Immunostaining of 3-month (left) and 8-month (right) MGEOs shows voltage-gated potassium channel K_v_3.1 (green) co-expression with MEF2C+ (red) at 8 months. Dashed boxes shown at higher-magnification on the right. (I) Confocal images of immunostaining shows PV (green) co-expressed with the perineuronal net marker WFA (red) in 6-month MGEOs. Dashed boxed area shown in higher magnification on the right. (J) Confocal imaging of PV basket cell-specific AAV1-BiPVe3-dTom virus expression in live-imaged 6-month MGEOs reveals complex morphologies of putative PV basket cells.

MGEOs at 1 and 3 months exhibited robust expression of MEF2C (**Figure 3A, G**), an early marker of PV-fated cortical interneurons^51,52^. Immunostaining of 3-month MGEOs further revealed a small number of PV-immunoreactive cells (2.2% ± 0.63 of DAPI area, **Figure 3E, F**). At 4 months, however, PV immunolabeling robustly increased (32.8% ±7.52) and was maintained at 6 (31.3% ± 7.06) and 8 months (32.5% ± 5.75) (**Figure 3E-G**). PV fast-spiking cortical interneurons express K_v_3 family potassium channels, predominantly K_v_3.1 and K_v_3.2^53^. Although expression of *KCNC1*, which encodes K_v_3.1, was already present in 1-month MGEOs (**Figure 3A**), there was minimal K_v_3.1 immunolabeling in 3-month MGEOs **(Figure 3H**). However, K_v_3.1 immunolabeling increased markedly in MGEOs by 8 months, and was seen predominantly in MEF2C+ cells (**Figure 3H**). In 6-month MGEOs, we observed PV-immunoreactive cells co-expressing Wisteria floribunda agglutinin (WFA), a marker for perineuronal nets that surround fast-spiking interneurons (**Figure 3I**)^54,55^. Viral labeling using an AAV1 PV basket cell-specific enhancer-driven tdTomato virus (AAV1-BiPVe3-dTom)^56^ confirmed the presence of putative PV basket cells with complex morphologies in 6-month MGEOs (**Figure 3J**). Together, these data indicate that MGEOs robustly produce PV+ cortical interneurons within 4 months in culture that exhibit molecular and structural features consistent with fast-spiking activity.

### MGEO transcriptome closely resembles that of the developing human MGE

To assess the fidelity of the MGEO model in recapitulating the transcriptomic features of the developing human MGE, we compared scRNAseq data of 90 DIV MGEOs to a published human fetal GE scRNA-seq dataset^11^. The human fetal GE dataset was visualized by UMAP plot using cell cluster annotations originally assigned in the dataset and then integrated with our MGEO dataset, and the relationships analyzed using UMAP and PCA plots (**Figure 4A-D**). The analysis shows a close overlap of MGEOs with the human fetal MGE cluster, suggesting that the transcriptome of the organoids closely mirrors that of the human fetal MGE. We next conducted Pearson correlation analysis to compare the neuronal clusters in the MGEOs with clusters from the human fetal GE dataset. A high degree of similarity between the corresponding neuronal subtypes in both datasets was revealed (**Figure 4E**), reinforcing the fidelity of MGEOs as a model of the human MGE.

**Figure 4.**
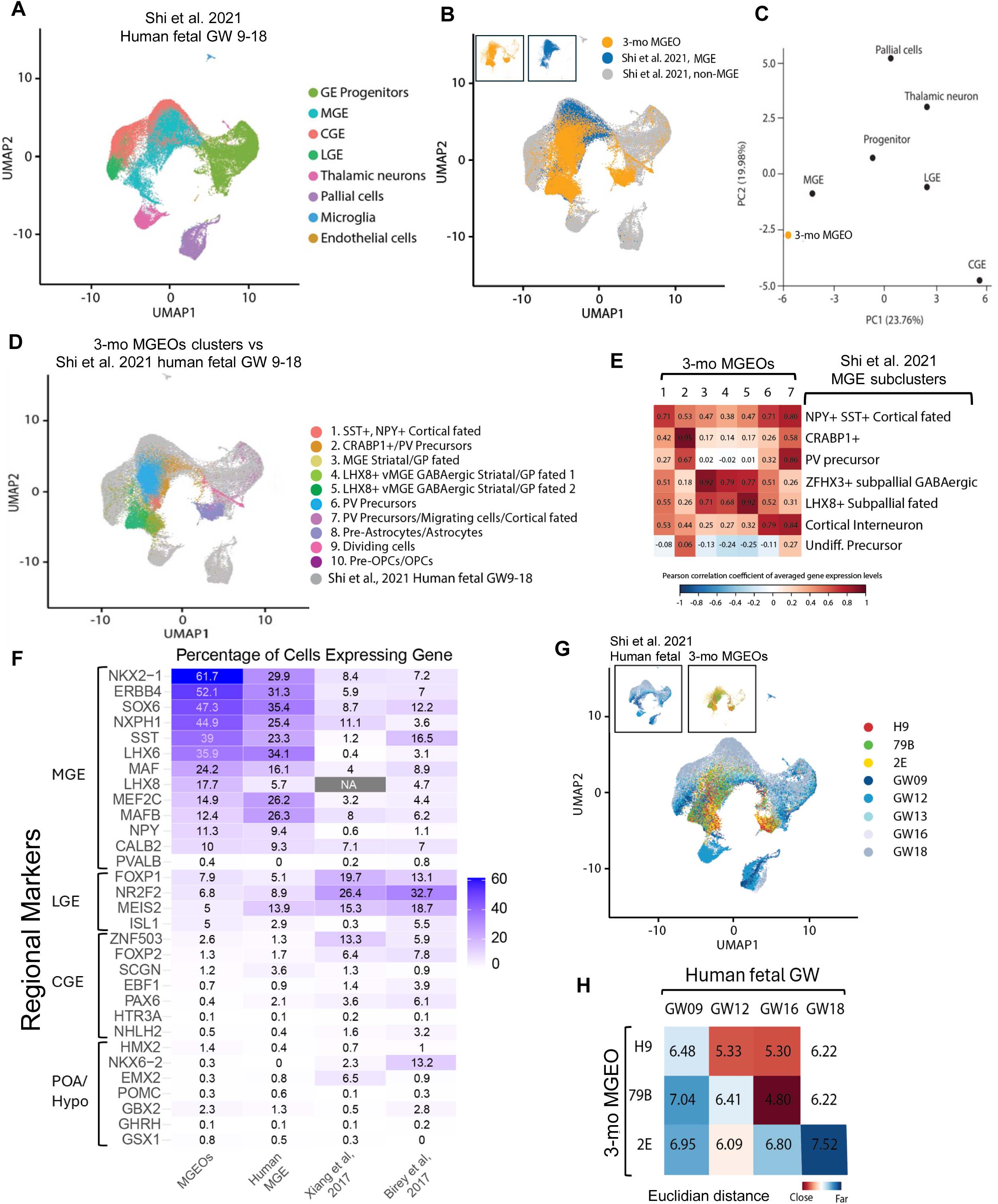
Comparison of MGEO and developing human subpallium transcriptomes (A) UMAP plot of human GW9-18 scRNA-seq data from Shi et al., 2021, with color-coded regional and cell type clustering. (B) Integrated UMAP plot of scRNAseq data from 3-month MGEOs (orange) compared with data from panel (A). Human fetal MGE cluster: Blue; non-MGE clusters: grey. Insets show the MGEO and human MGE clusters individually. (C) Principal component analysis (PCA) of 3-month MGEO scRNA-seq data (orange dot) versus human fetal data from (A) shows that MGEOs map closest to the human MGE cluster. (D) UMAP showing 3-month MGEO cell type clusters integrated with human GW9-18 fetal GEs (Shi et al., 2021; in gray). MGEO cell clustering and colors as in Figure 2D. (E) Heat map of Pearson correlation coefficients comparing 3-month MGEO clusters 1-7 (columns, numbered as in panel D) with Shi et al. 2021 human fetal MGE subclusters (rows). (F) Heat map comparing the average percentage of cells expressing selected gene markers for MGE, LGE, CGE, and preoptic area/hypothalamus (POA/Hypo) across scRNA-seq datasets from 3-month MGEOs, human fetal MGE (Shi et al., 2021; gestational weeks 9–18), and other subpallial organoids of similar timepoints (Xiang et al., 2017; Birey et al., 2017). (G) Integrated UMAP of 3-month MGEO and human fetal GE scRNA-seq data, as shown in panels B and D. Developmental age clusters from Shi et al. are depicted in shades of blue, while 3-month MGEO cells are colored by cell line (red: H9; green: 79B; yellow: 2E). Insets display human fetal cells (left) and MGEO cells (right) separately. (H) Heatmap of PCA Euclidean distances comparing each 3-month MGEO cell line to human fetal gestational developmental ages (GW 9, 12, 16, and 18) shows 3-month MGEOs are, on average, closest to human developmental GW12-16.

We then compared gene expression between the MGEOs, human MGE, and two published subpallial organoid models^11,32,33^ by analyzing the percentage of cells expressing genes specific to various regions of the developing subpallium (**Figure 4F**). The results demonstrated that MGEOs exhibit strong expression of MGE-specific markers, with minimal off-target expression of genes associated with other brain regions including LGE, CGE, POA, and hypothalamus. In contrast, the two multi-rosette subpallial organoid protocols^57,58^ yielded regionally mixed expression patterns and limited MGE specificity. Furthermore, we found that the MGEOs produced much larger percentages of GABAergic neurons and MGE-specific GABAergic neuron subtypes (**Figure S3C**). Several cell populations in 3-month MGEOs expressed markers of cortical PV precursors, including *LHX6, MEF2C, MAFB, ERBB4, ETV1*, or markers of subpallial destined PV precursors, with co-expression of LHX8, TAC1, and CRABP1 (**Figure S3D**). Detailed analyses of gene expression patterns of GABAergic neuron subtypes in the MGEOs further revealed that many cells co-expressed *SST*, including *SST*+*/CHODL*+, *SST*+*/CALB2*+, and *SST*+/*NPY*+ combinations, known *SST* subtypes generated in the MGE^59,60^. Notably, *SST, CHODL, CALB2*, and *NPY* were largely not co-expressed in PV+ (*PVALB*) cells, consistent with prior studies showing that PV⁺ and SST⁺ interneurons, though both derived from the MGE, represent virtually non-overlapping cell populations^61^ (**Figure S3E)**. Integration of MGEO data from multiple cell lines with human data across various gestational weeks showed that 3-month MGEO clusters closely aligned with those of the human MGE at similar stages of development, demonstrating the robustness of the MGEO model across cell lines (**Figure 4G**). To further investigate the developmental trajectory of the MGEOs, we computed Euclidean distances between MGEOs and the human dataset^11^ at different gestational weeks (**Figure 4H**). The heatmap revealed that the 3-month MGEOs most closely resemble human MGE tissue from gestational weeks 12-16. Together, these data show that the MGEO model captures key features of MGE cellular diversity and accurately recapitulates the transcriptomic signature of the developing human MGE.

### MGEOs recapitulate cortical interneuron migration and structural integration in assembloids

Given that MGE-derived cortical interneurons are highly migratory and possess migratory features that differentiate them from radially migrating cortical neurons^62,63^, we characterized migratory properties of interneuron progenitors in MGEOs alone and in assembloids after fusion with COs. MGEOs at different time points were first plated on extracellular matrix (ECM)-coated plates (**Figure 5A).** Day 21 was the earliest time at which neuron-specific migration appeared without the spread of proliferative cells (**Figure 5B-C**). Neuronal migration onto the ECM was robust and rapid, as seen by comparing images at 0-, 12-and 24-hours post-plating (**Figure 5B’, Video S1**). Cells that migrated away from the MGEOs co-expressed MAP2ab and LHX6, supporting their neuronal and MGE lineage identity (**Figure 5C**).

**Figure 5:**
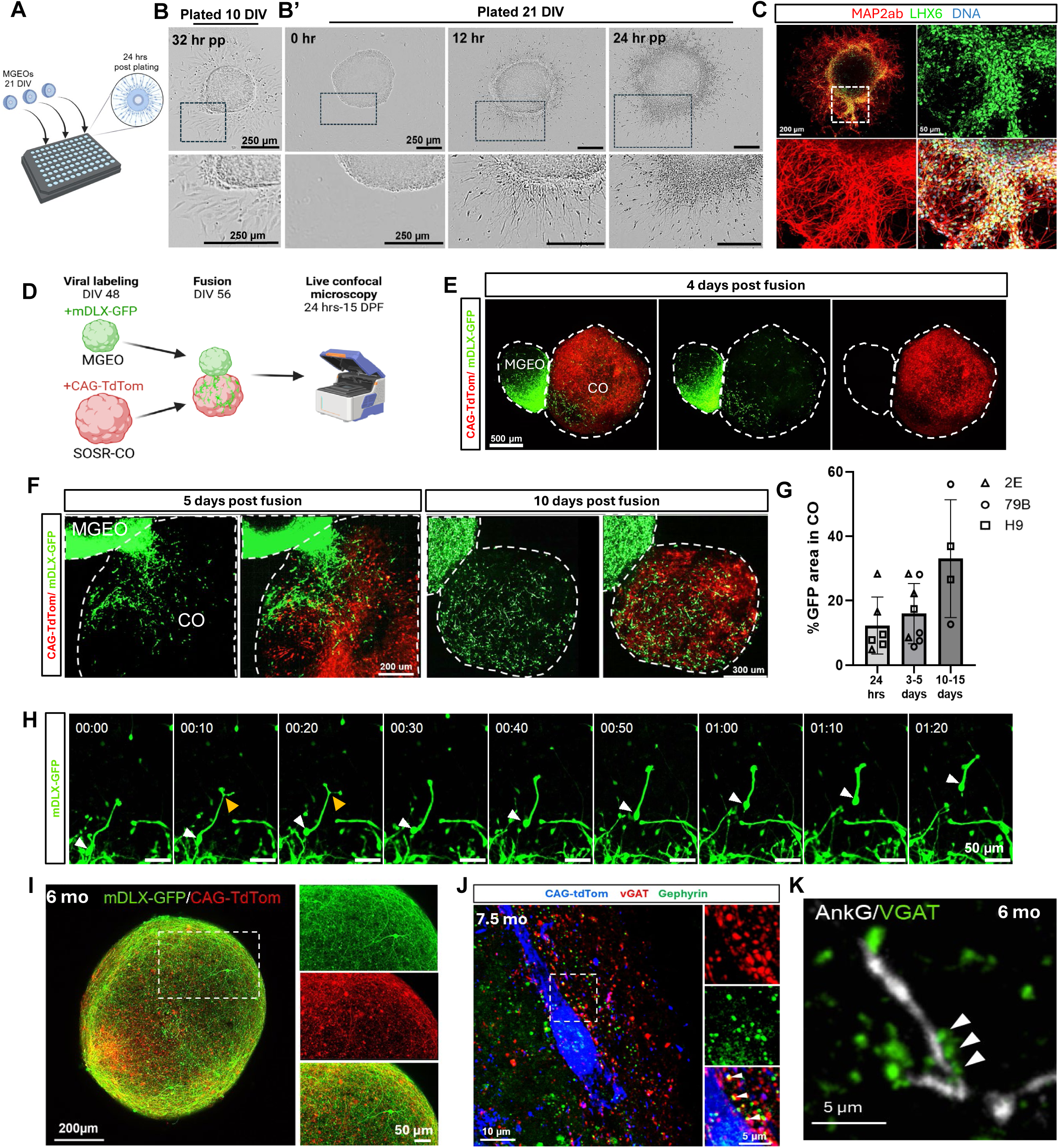
MGEO-derived interneuron progenitors are highly migratory (A) Diagram of 2D migration assay. (B) Images of MGEOs at 32 hours post-plating (hr pp) at 10 DIV, and (B’) 0, 12, and 24 hr pp at 21 DIV on Geltrex. Also see **supplemental video S1**. (C) Confocal images of a MGEO at 3 days pp immunolabeled for MAP2ab (red) and LHX6 (green) showing co-expression in MGE-derived neurons. Bisbenzimide (DNA) in blue. Boxed region of upper left image is shown at higher magnification in the other panels. (D) Schematic of assembloid protocol, involving viral labeling at 48 DIV, fusion at 56 DIV, and subsequent live confocal microscopy from 1-15 days post fusion (DPF). (E-F) Images from representative MGEO-CO assembloids at 4 (E) or 5 and 10 (F) DPF. mDLX-GFP (green) virus labels GABAergic cells originating from the MGEO, whereas tdTomato (red) labels CO progenitors and putative excitatory neurons. Also see **supplementary video S2**. (G) Quantification of mDLx-GFP+ virally labeled MGEO interneuron migration into COs at 1, 3-5, and 10-15 DPF. Percentage area of GFP expression in the CO organoid ± standard deviation (SD) is shown. Points represent single organoid replicates (n≥4), with the point shape denoting the cell line used (triangle: 2E, circle: 79B, diamond: H9). (H) Timelapse live imaging of a mDLX-GFP-labeled MGEO interneuron within the CO region of a MGEO-CO assembloid shows characteristic interneuron leading edge branching movement patterns. Time is listed as hr:min. The white arrow denotes the neuronal soma, and the yellow arrowhead points to branching of the leading process. Also see **supplementary video S3**. (I) Confocal image of an MGEO-CO assembloid at 6 months, immunolabeled with mDLX-GFP (green) and tdTomato (red) as in D-F, shows full fusion into a single spherical structure. The boxed area at left is expanded in the right panels. (J) Confocal image of a 7.5 month MGEO:CO assembloid immunolabeled for GABAergic post-and pre-synaptic markers Gephyrin (green) and vGAT (red), respectively, on a CAG-tdTom CO neuron (pseudocolored blue). Boxed area in the left panel is shown at higher magnification at right, with white arrowheads in the merged image showing closely apposed vGAT+ and Gephyrin+ puncta. (K) Immunolabeling of a 6-month MGEO:CO assembloid shows VGAT puncta (green) localized along an Ankyrin G (AnkG)-positive axon initial segment (white).

Cortical interneurons migrate tangentially from subpallial regions to the cortex and are directionally guided by attractive and repulsive cues expressed in the cortex and subpallium^64,65^. For example, binding of the α-chemokine SDF-1 (CXCL12) to the G-protein-coupled CXCR4 receptor in migrating interneurons facilities tangential migration (**Figure S4A**)^66,67^. To test whether this chemokine signaling influences MGEO-derived interneurons, we first confirmed that *CXCR4*, *SDF-1*, and *CXCR7*, another chemokine receptor involved in interneuron migration, were expressed in our 1-month MGEO scRNA-seq dataset (**Figure S4B**). Next, we plated MGEOs on Geltrex at day 30 and assayed for cell migration over 2 days while exposing them to 100 nM of the CXCR4 inhibitor AMD3100 for 8 days (Chronic; 6 days before and 2 days after plating) or 2 days (Acute; after plating) (**Figure S4C**). We measured the area of migrated neurons outside the organoids at 48 hours post plating. Although the average migrated neuron area from MGEOs in the Acute treatment group was not statistically different from vehicle-treated controls, MGEOs chronically treated with AMD3100 showed significantly reduced neuronal migration vs control (**Figure S4D, E**). The reduced migration of MGEO-derived putative interneurons with CXCR4 antagonist treatment suggests that cortical interneuron migration from MGEOs is influenced by mechanisms similar to those seen *in vivo*.

To examine migration using an assay that more faithfully recapitulates *in vivo* cortical interneuron migration, we fused MGEOs with dorsally patterned SOSR-COs (hereafter referred to as “COs”)^38^ at 56-58 DIV to generate assembloids. Prior to fusion, MGEOs and COs were virally labeled with mDLX-GFP^68^, and CAG-tdTomato, respectively (**Figure 5D**). We observed robust migration of mDLX-GFP+ interneurons that began several hours after fusion and continued rapidly throughout the first two weeks post-fusion (**Figure 5E-G, Video S2 and data not shown**). The rapid migration of mDLX-GFP+ interneurons was consistent across cell lines and experimental replicates (**Figure 6G**). Migrating interneurons displayed properties indicative of cortical interneuron migration, including somal translocation and leading-edge branching in MGEO-CO assembloids **(Figure 6H, Video S3**) as well as in 2D assays described above (**Video S1**). Notably, MGEO-CO fusions treated with AMD3100 1-day post fusion trended toward reduced mDLX-GFP+ interneuron migration into the COs versus controls at 48-hours post fusion (**Figure S4F**).

**Figure 6.**
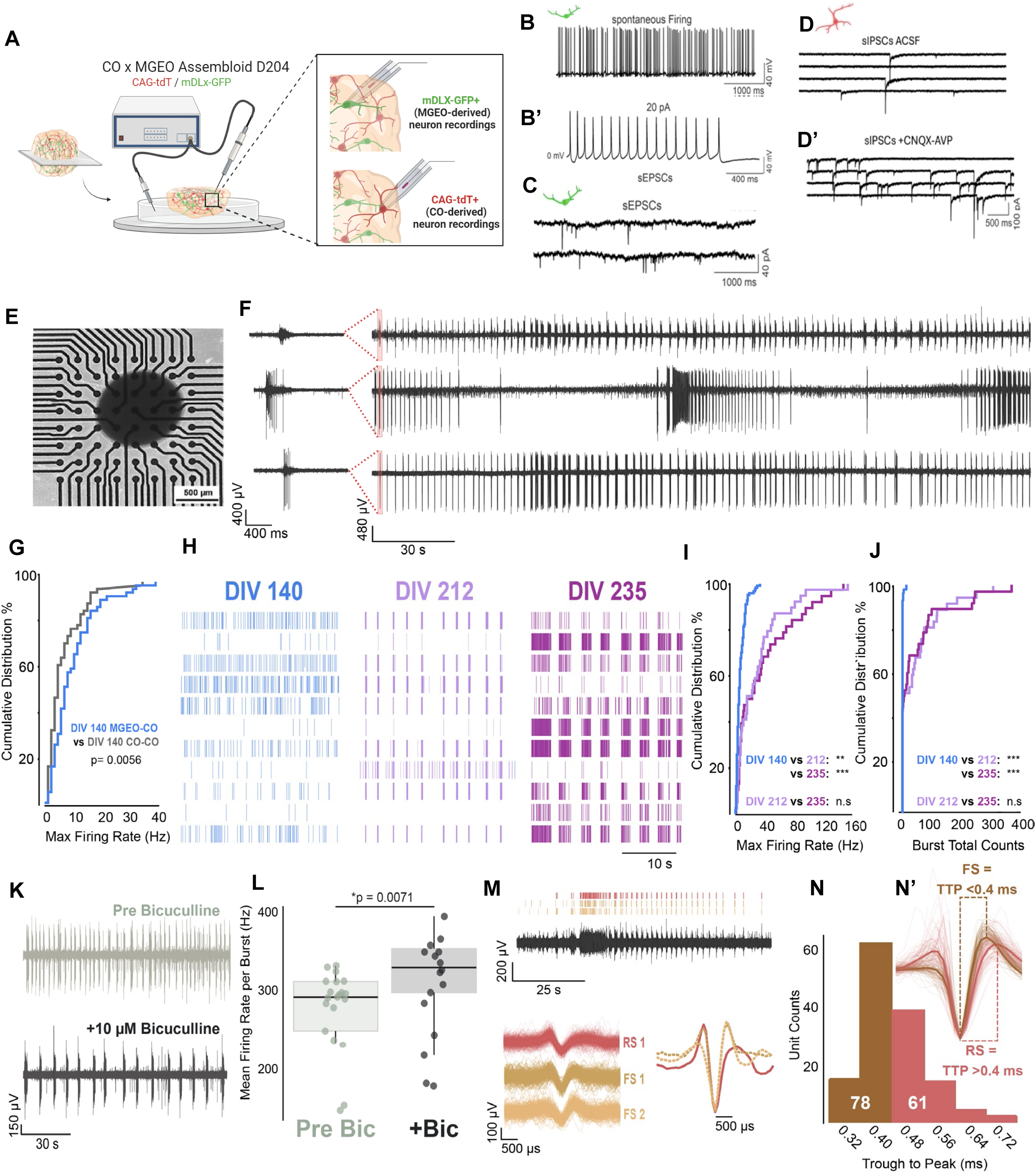
MGEO-CO assembloids form functional synapses, display increasingly complex network activity and exhibit fast-spiking units (A) Diagram of assembloid acute slice whole-cell patch clamp recordings. (B) Representative traces showing spontaneous (B) and evoked (B’) repetitive firing of mDLX-GFP+ (MGEO-derived) neurons in MGEO-CO assembloid slices at 204 DIV. AP firing was evoked from RMP by injection of 1500 ms pulse currents (20 pA). (C) Representative trace of spontaneous excitatory postsynaptic currents (sEPSCs) recorded from an mDLX-GFP+ neuron within an assembloid slice at 204 DIV (holding potential-70 mV). (D) Representative trace of spontaneous inhibitory postsynaptic currents (sIPSCs) recorded from a CAG-tdT+ (CO-derived) neuron in the same slice as (C). (D’) sIPSC were recorded at a holding potential of-70 mV in the presence of 6-cyano-7-nitroquinoxaline-2,3-dione (CNQX, 10 µM) and amino-5-phosphonopentanoic acid (APV, 50 - 100 µM). (E) Brightfield image of a 6-month MGEO-CO assembloid acutely plated on an MEA plate for recording. (F) Representative MEA voltage traces from 3 electrodes taken from 2 representative MGE-CO assembloid recordings shown over 2 s (left) and 5 min (right) windows. (G)Cumulative distribution of maximum firing rates between DIV-140 CO-CO (gray) and MGEO-CO (light blue) assembloids. n=3 each from 2 cell lines with 62 putative units/assembloid type. Two-tailed Kolmogorov-Smirnov test. (H) Representative raster plots of putative units in MGEO-CO assembloids at different maturation stages: DIV-140 (n=3), DIV-212 (n=4-6), and DIV-235 (n=4-6) from 2 cell lines. (I-J) Cumulative distribution of maximum firing rates (I) and burst counts (J) across maturation stages (DIV-140, DIV-212, and DIV-235) using same samples as in H. Two-tailed Kolmogorov-Smirnov test, DIV 140 vs 212: (I) KS=0.5010, **, P=2.3247e-07, (J) KS=0.5062, ***, P=1.6290 x10^-7^. DIV 140 vs 235: (I) KS Stat=0.4597, ***, P=4.1009e-06, (J) KS Stat=0.5123, ***, P=1.5173 x10^-7^. DIV 212 vs 235: (I) KS Stat=0.1350, n.s., P=8.1858e-01, (J) KS=0.1404, n.s., P=0.77593. N=38-62 putative units/3-5 assembloids per timepoint. (K) Electrode-matched representative voltage traces from a 5-min recording before (gray, top) and after (black, bottom) application of 10 µM bicuculline. (L) Firing rate per burst of putative units. Boxplots display the median, 25th and 75th percentiles (box edges), whiskers extending to the most extreme data points, and individual outliers plotted separately. Two-tailed Mann-Whitney U=113, Pre Bic treatment: N=9-11 putative units/3 MGEO-COs, During Bic treatment: 8-10 putative units/2 MGEO-COs. (M) Representative voltage trace with corresponding raster plots of three putative units recorded during a network burst. Red represents a regular-spiking unit (RS1); brown (FS1) and gold (FS2) represent fast-spiking (FS) units. (N) Quantification from panel (M): histogram distribution of putative FS vs. RS units pooled from n=10 assembloids across ≥2 independent differentiations each from 2 cell lines. (N’) representative voltage traces of putative RS and FS units aligned to the trough. The mean trough-normalized waveforms of each unit are overlaid, showing trough to peak (TTP) measurements and non-overlapping RS and FS waveforms. MEA data pooled from 2-4 independent differentiations.

After ∼5 months post fusion, MGEOs fully merged with COs and the entire assembloid became a single spherical structure (**Figure 5I**). mDLX-GFP+ interneurons within assembloids co-expressed GABA (**Figure S5A**) and exhibited robust process outgrowth with both bipolar and multipolar interneuron morphologies (**Figure S5B-C**).

Immunostaining of assembloids at 9 months revealed robust expression of PV and Kv3.1, especially in the MGEO periphery (**Figure S5D-F**), and the formation of inhibitory synapses with vGAT+ puncta closely apposed to Gephyrin+ puncta on the somas of putative cortical excitatory neurons marked by CAG-tdTomato (**Figure 5J**). vGAT+ puncta were also identified along Ankyrin G immunolabeled axon initial segments (**Figure 5K**), indicative of inhibitory axo-axonic synapses. These data provide structural evidence for the maturation of MGEO-derived interneurons and their synaptic integration with excitatory neurons in assembloids.

### MGEO-CO assembloids display functional integration of interneurons, complex activity and fast-spiking neurons

To provide evidence for functional synapses and examine neuronal activity, we next performed whole-cell patch-clamp and multielectrode array (MEA) recordings from MGEO-CO assembloids. Action potential (AP) firing and spontaneous post-synaptic currents in both excitatory and inhibitory neurons were examined using whole-cell recordings of slices made from 6-month assembloids (**Figure 6A-D**). We observed spontaneous AP firing in 50% (9 of 18 cells) of recorded neurons, and an additional 17% (3 of 18 cells) displayed repetitive AP firing upon depolarizing current injection (**Figure 6B, B’**). We also found that glutamatergic and GABAergic neurons formed functional connections, with recordings of mDLX-GFP-labeled interneurons displaying spontaneous excitatory post-synaptic currents (sEPSCs; **Figure 7C**) and CAG-tdTomato-labeled, putative excitatory neurons receiving spontaneous inhibitory post-synaptic currents (sIPSCs; **Figure 7D**). sIPSCs were also recorded after co-application of the AMPA and NMDA receptor antagonists CNQX and APV to isolate GABAergic synapses (**Figure 7D’**).

**Figure 7.**
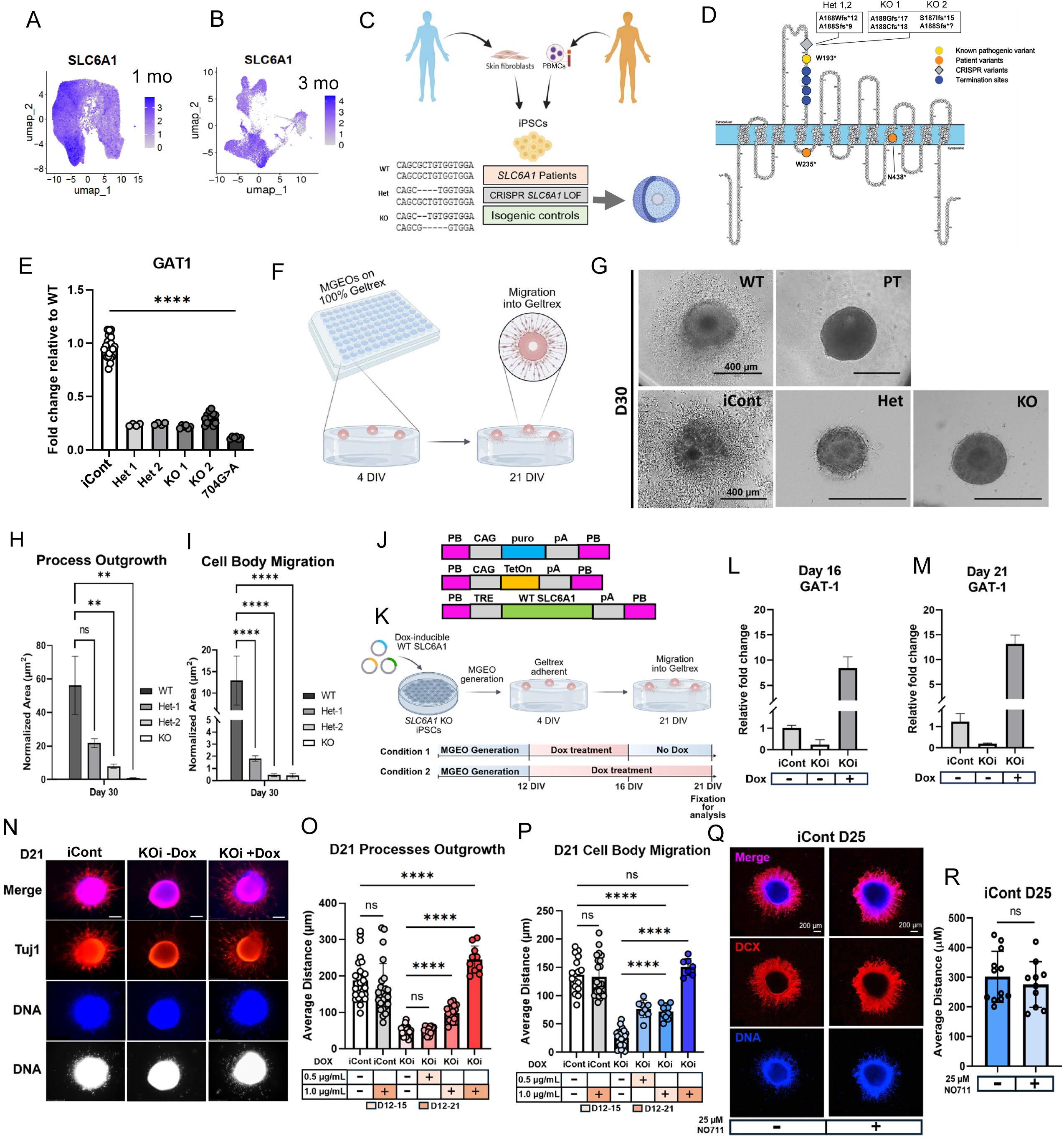
*SLC6A1* variants cause migratory deficits in developing human interneurons (A, B) UMAP plots showing *SLC6A1* expression in 1-month (A) and 3-month (B) MGEOs. (C) Schematic describing the generation of CRISPR gene edited and patient-derived iPSC lines. (D) Representation of GAT1 protein structure showing location of CRISPR generated indels, predicted truncation sites, and patient variants. (E) Analysis of RT-qPCR fold change in GAT1 mRNA expression levels in *SLC6A1* variant lines relative to isogenic control (iCont). (F) Schematic representation of MGEO migration assay. (G) Brightfield images of cell migration from MGEOs into the surrounding extracellular matrix at day (D) 30 in vitro. (H, I) Quantification of MGEO process outgrowth area (H) and cell body migration area outside of MGEOs (I) in WT, 2 Het and KO groups. (J, K) PiggyBac constructs (J) and experimental workflow (K) for doxycycline (Dox)-inducible *SLC6A1* rescue experiments. (L, M) Analysis of GAT1 mRNA expression with (+) or without (-) Dox in DIV 16 (L) and DIV 21 (M) MGEOs derived from *SLC6A1* Dox inducible KO lines (KOi) and untreated isogenic control (iCont). (N) Confocal images showing migrating cells from iCont and KOi DIV 21 MGEOs embedded in ECM and immunolabled for TuJ1 (red) to identify processes and bisbenzimide labeling (blue) for DNA, which is also pseudocolored white for better visualization of migrated cells. (O, P) Quantification of average distance of process outgrowth (O) or cell migration (P) in iCont and KOi MGEOs without Dox (-) and with two different DOX concentrations (+) for two experimental durations (D12-15) and (D12-21). Each data point represents individual organoids from 3 separate experiments. (Q) Confocal images of doublecortin (DCX)-immunolabeling (red) shows cell migration and process outgrowth from iCont DIV 25 MGEOs 4 days after plating on ECM with (+) or without (-) the addition of the GAT1 blocker NO711. Bisbenzimide DNA labeling in blue. (R) Quantification of average cell body migration distance in the presence or absence of the GAT1 blocker NO711. ns, non-significant.

Most published MEA recordings from brain organoids are conducted with the organoids chronically cultured on MEA plates, leading to the migration of many neurons onto the plate surface^69^. This approach disrupts the 3D cytoarchitecture and records activity from both 2D and 3D networks that may not faithfully replicate activity exclusively derived from 3D structures. To overcome this limitation, we transiently plated MGEO-CO assembloids grown in suspension onto MEA plates for recordings (**Figure 6E**), before placing them back in suspension culture, allowing for recordings made exclusively from 3D networks and recordings from the same organoid at later timepoints (see **Methods**). Inspection of the raw voltage time series revealed large-amplitude voltage fluctuations, the presence of both single-and multi-unit APs, and bursts of APs (recordings made at 212 and 235 DIV) (**Figure 6F**). We hypothesized that MGEO-CO assembloids would contain a significant number of MGE-derived high-firing-rate interneurons, as compared to CO-CO assembloids without interneurons. Consistent with this prediction, 140 DIV CO-CO assembloids exhibited a larger fraction of putative units with low firing rates relative to MGEO-CO assembloids (**Figure 6G**). This finding suggests that some MGE-derived interneurons in MGEO-CO assembloids include fast-spiking interneurons, aligning with the observed increase in high-firing-rate units in MGEO-CO assembloids.

Raster plots of MGEO-CO assembloids recorded at DIV 140, 212, and 235 illustrate the expected emergence of network complexity and synchronicity as a function of organoid maturation (**Figure 6H**). Cumulative distribution analyses of maximum firing rates across different maturation stages revealed that firing rates at DIV 140 are significantly lower compared to those at DIV 212 and 235, while no significant difference was observed between DIV 212 and 235 (**Figure 6I**). Similarly, the cumulative distribution of burst numbers increased over time, as DIV 140 assembloids exhibited significantly fewer bursts compared to the later time points **(Figure 6J**). No significant difference in number of bursts was observed for MGEO-CO assembloids between DIV 212 and 235 **(Figure 6J**); however, complex bursting patterns were observed in MGEO-CO assembloids at DIV 235 (**Figure S6**). Collectively, these data suggest that as MGEO-CO assembloids mature, both firing rates and network bursting activity increase and shift to more complex bursting patterns.

To further confirm the presence of functional interneurons in MGEO-CO assembloids, we applied 10 µM bicuculline, a GABA-A receptor antagonist, to assess the role of GABAergic transmission in regulating network activity. Electrode-matched representative voltage traces from a 5-minute recording before and after bicuculline application showed clear alterations in firing rates with bicuculline exposure (**Figure 6K**). Quantification of this effect confirmed that blocking GABAergic transmission increases firing rates within network bursts (**Figure 6L**). In addition, we classified putative units from extracellular recordings to distinguish between fast-spiking (FS) and regular-spiking (RS) neurons using spike sorting and measurements of trough to peak (TTP) AP duration (**Figure 6M, N**).The majority of putative units in MGEO-CO assembloids were FS units, further supporting the presence of MGE-derived FS interneurons. These results provide direct evidence that inhibitory neurons in MGEO-CO assembloids are functionally integrated and that FS units are effectively generated in MGEO-CO assembloids.

### *SLC6A1* LOF results in cortical interneuron cell-autonomous migratory deficits

We next looked at the expression of a select set of genes associated with human neurodevelopmental disorders in our MGEO scRNA-seq data. We compared gene expression at 1-and 3-month timepoints and across different cell type clusters (progenitors/glia, cortical interneurons, and subpallial neurons). We found that many of these genes were expressed in our MGEOs (**Fig. S7**), offering the opportunity for human cell-based disease modeling. Notably, we found that *SLC6A1*, which encodes the plasma membrane GABA transporter-1 (GAT1) that is highly expressed in cortical interneurons^70–72^, showed strong expression in the MGEO cortical interneuron clusters at 1 and 3 months and its expression was especially upregulated in cortical PV precursor clusters at 3 months (**Figs. 7A and S7**). Pathogenic *SLC6A1* variants resulting in haploinsufficiency are linked to myoclonic atonic epilepsy (MAE, MIM# 616421)^73–75^, and this gene is also implicated in autism and other neuropsychiatric disorders^76^.

To model MAE using MGEOs and explore potential effects of *SLC6A1* loss of function on interneuron development, we used CRISPR/Cas9 editing to generate loss-of-function variants and also reprogrammed *SLC6A1*-linked MAE patient cells into iPSCs (see Methods). We performed dual fibroblast reprogramming and *SLC6A1* CRISPR/Cas9 gene editing^77^ to create out-of-frame insertions/deletions in the *SLC6A1* gene, near a known W193* truncation variant linked to MAE with mild autistic features^73^. Two heterozygous (Het 1, 2), two compound heterozygous (KO 1, 2), and isogenic control (iCont) lines were generated. We also reprogrammed PBMCs from both male and female patients with *SLC6A1*-linked MAE (PT 1, 2) and from an unrelated healthy control (WT) (**Figure 7C,D**). Together these variants spanned various regions of the GAT1 protein including extracellular, intracellular, and transmembrane domains (**Figure 7D**). GAT1 mRNA reduction was confirmed in KO, HET and PT lines using RT-qPCR, with GAT1 expression between 5-40% of WT levels (**Figure 7E**).

The migration of GE interneurons to the cortex parallels that of GAT1 expression during development^71^. Additionally, ambient GABA levels have been shown to alter interneuron migration through non-synaptic interactions at GABA receptors^78–80^. To evaluate the effects of *SLC6A1* LOF on cortical interneuron migration, MGEOs from 1 each of PT, KO, HET, iCon and WT groups were left embedded in thick extracellular matrix (100% Geltrex; part of the single-rosette MGEO protocol) and both somal migration distance and process extension into the matrix were quantified at DIV 30 (**Figure 7F-I**). We observed a significant reduction in cell migration and process extension from PT-derived, Het, and KO MGEOs compared to iCont and WT. This finding was unexpected as migration deficits have not been reported in *SLC6A1* mouse models.

To determine if *SLC6A1* loss was indeed responsible for these migration deficits, we conditionally restored *SLC6A1* expression using PiggyBac transposase to stably integrate a doxycycline (Dox) inducible, codon-optimized, wildtype *SLC6A1* cassette into the *SLC6A1* KO line (KOi) (**Fig 7J**). DIV 12 MGEOs embedded on top of thick ECM were exposed to either 0.5 or 1.0 ug/mL Dox in the culture media for either 3 or 9 consecutive days (**Fig 7K**). Controls included iCont MGEOs with and without Dox, vehicle exposure of KOi MGEOs, and Dox exposure of MGEOs without the overexpression cassette. Increases in WT GAT1 mRNA in the KOi MGEOs were confirmed via RT-qPCR at DIV 16 and 21 (**Fig 7L, M**). Migration distance of cell bodies and extension distance of processes were assessed at DIV 21 after labeling with Hoechst and immunostaining with TuJ1 antibody, respectively. We found that WT *SLC6A1* overexpression in the KOi MGEOs rescued the migration and process outgrowth deficits in a Dox dose and duration dependent manner (**Fig 7N-P**). We next asked whether inhibiting GAT1 transport activity in control MGEOs would impair interneuron migration. Interestingly, blockade of GAT1 from DIV 12-21 using the GAT1-specific blocker NO711 (also known as NNC-711) (25 µM) did not inhibit migration from iCont MGEOs (**Fig 7Q-R**). These findings indicate that GAT1 expression is necessary for the appropriate migration of MGE-derived interneurons, and that GAT1 may play a non-canonical role in early interneuron migration that is separate from its function as a GABA transporter.

To assess the impact of *SLC6A1* loss of function on interneuron migration in an organoid system, we generated assembloids of mDLX-GFP labeled MGEOs and CAG-TdTom labeled COs from iCont, WT, Het, and PT-derived lines (**Figure 8A**), the latter two groups modeling disease-associated haploinsufficiency. Consistent with our findings above, we observed significantly reduced migration of interneurons into the CO region in all Het and patient groups compared to iCont and WT (**Figure 8A, B**). Mutant mDLX-GFP+ cells appeared to accumulate at the cortical organoid junction with very few cells able to cross this boundary. No significant differences were seen between iCont and WT groups. We further assessed if the migration deficit reflected cell autonomous or non-cell autonomous effects by creating “mixed and matched” assembloids using WT or PT COs fused with either WT or PT MGEOs (**Figure 8C**). Defective mDLX-GFP+ cell migration appeared in all conditions containing PT MGEOs, regardless of the CO genotype. A migration deficit was not observed in any condition containing WT MGEOs irrespective of the CO genotype (**Figure 8D)**. These results suggest that GAT1 haploinsufficiency during early development causes a cell-autonomous impairment in MGE cortical interneuron migration.

**Figure 8.**
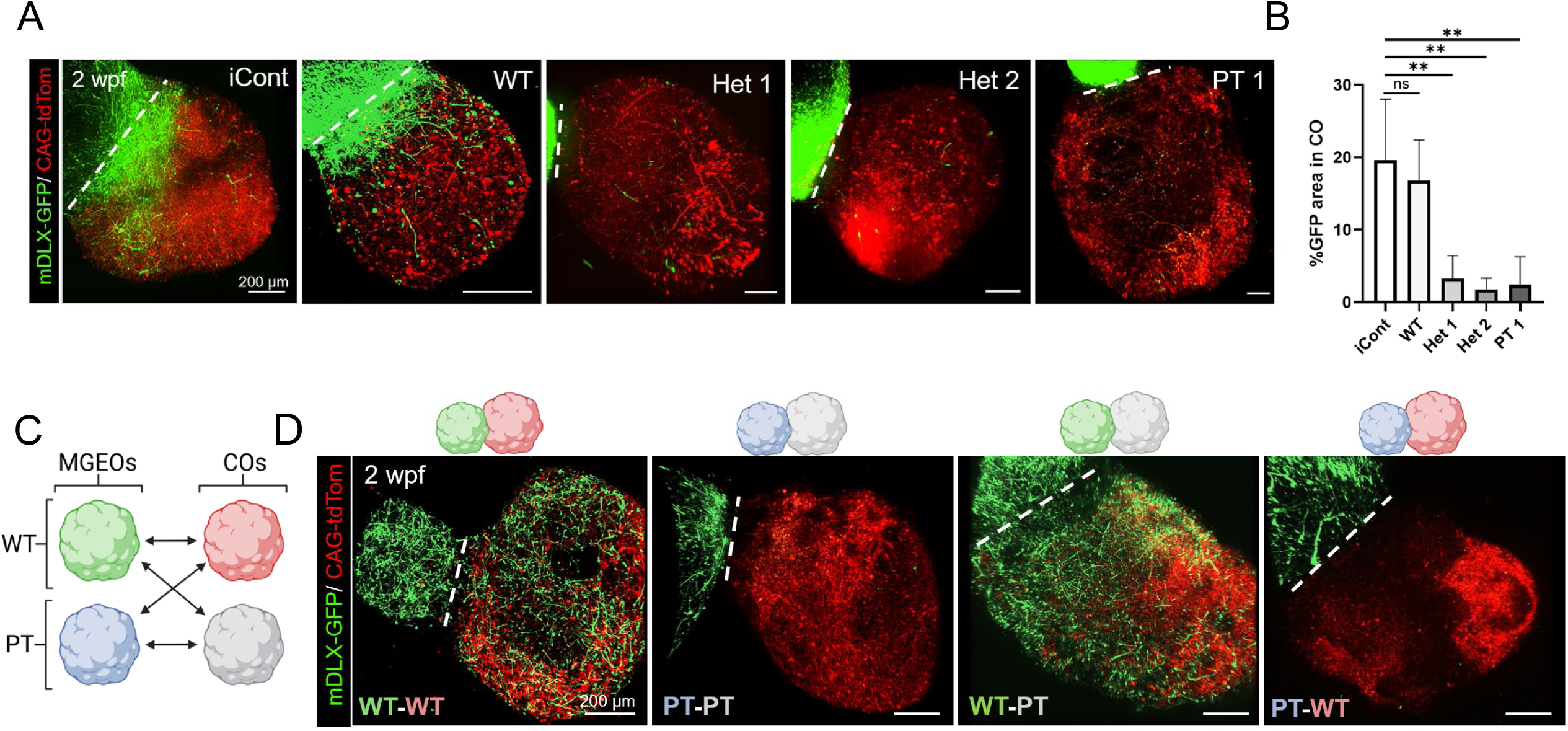
SLC6A1 migratory deficits are cell autonomous in MGEO-CO assembloids (A) Representative live confocal images of MGEO-CO assembloids at 2 weeks post fusion (wpf) and derived from isogenic control (iCont), wildtype (WT), two *SLC6A1* heterozygous KO (Het 1, 2) and *SLC6A1* patient (PT1) lines. mDLX-GFP (green) virus labels GABAergic cells originating from the MGEO, and tdTomato (red) labels CO progenitors and putative excitatory neurons. Note decreased MGEO cell migration into COs in HET and PT assembloids. Images are max projections of 150 µm thick assembloid sections. The dashed lines denote the boundary between MGEO and CO. (B) Quantification the average percentage area of mDLX-GFP expression in the CO organoid area ± SD is shown for 6-9 assembloids from 2 separate experimental replicates per cell line for the groups from (A). **, p<0,0023; ns, non-significant. (C) Schematic representation of assembloid variations in “mix and match” experiments. (D) Live confocal images of genotype mixed and matched MGEO-CO assembloids at 2 wpf derived from wildtype (WT) and patient (PT) lines. MGEO migration into assembloids is markedly decreased in assembloids with PT MGEOs, regardless of whether the CO is from the PT or WT, but WT MGEOs show intact migration into PT COs. Images are max projections of 150 µm thick assembloid sections. The dashed lines denote the boundary between MGEO and CO.

## Discussion

A critical challenge for studying human cortical development and brain disorders with human cellular models is the inability to generate late developing PV interneurons.

These cells arise from the embryonic MGE, and nearly all existing subpallial organoid models are very limited in generating MGE lineages, especially PV. Here we show the development of an MGE-specific subpallial organoid protocol, MGEOs, that robustly and consistently produces MGE lineages closely resembling the embryonic human MGE and gives rise to highly migratory interneuron progenitors that differentiate into PV and SST interneurons. These interneurons integrate with excitatory cortical neurons in assembloids that exhibit complex network activity and show a substantial proportion of fast-spiking neurons. We also demonstrate that *SLC6A1* haploinsufficiency linked to MAE disrupts MGEO-derived cortical interneuron migration in a cell autonomous manner.

We designed our small molecule manipulations to optimally pattern toward MGE, but starting with a hPSC monolayer was also probably advantageous. The even distribution of morphogens across all cells in the monolayer increases culture purity and better replicates the formation of the neural plate in human embryonic development. For multi-rosette organoid models that initiate with hPSC aggregation into embryoid bodies, the optimal timing and concentrations of patterning factor application may differ. Notably, a recent study described a multi-rosette subpallial organoid protocol with increased MGE specificity and the generation of some PV cortical interneurons in assembloids^118^. This group also reported the presence of PV reporter-labeled neurons with fast-spiking properties in their assembloids, although the firing rates were rarely above 60 Hz.

Another group enriched for MGE-like pallial interneurons in 2D culture and grafted them into mouse cortex ^29^. They successfully generated a large proportion of ERBB4+/SST-cells transcriptomically predicted to be of PV basket cell identity, but detected no *PVALB* mRNA up to 18 months post transplantation.^44^ In contrast, we observed robust immunoreactivity for PV in MGEOs (with or without fusion to hCOs) by 4 months, along with evidence of basket and axoaxonic subtypes and robust expression of K_v_3.1 and a perineuronal net marker, both of which are components of fast-spiking PV interneurons.

By MEA recordings, we identified fast-spiking neurons in assembloids with the top 20% at DIV 235 showing firing rates of 30-160 Hz, and the top 5 FS units above 80 Hz (**Figure 6I**). The detection of fast-spiking neurons in our assembloids likely reflects the capacity of MGEOs to generate PV cortical interneurons, along with the enrichment of interneurons at the assembloid periphery (see **Figure 3E**) in close juxtaposition to the MEA electrodes leading to a disproportionate identification of fast-spiking neurons. Our novel MEA recording approach, which allows for established 3D networks to remain intact as they mature rather than reorganizing into 2D structures^81^, also likely contributed. However, we did not have access to a PV reporter line for patch-clamp recordings to definitively show that the fast-spiking neurons are PV interneurons.

We focused on *SLC6A1* DEE to demonstrate the utility of MGEOs in disease modeling. The *SLC6A1* gene has been difficult to study in human model systems due to its limited endogenous expression in organoid and 2D neuronal models, likely reflecting the lack of MGE lineages in these models, particularly PV cortical interneurons. Enrichment of GAT1 in the cortex is found on presynaptic axon terminals of symmetrical synapses on the soma, and axon initial segment, characteristic of PV basket and axoaxonic cells^82,83^ which MGEOs readily produce. *SLC6A1* variant overexpression models provide insight into effects on transporter function^84–86^ but cannot address the potential role of endogenous GAT1 during early brain development. Animal models also do not fully recapitulate MAE disease phenotypes^87–89^. Here we identify a novel MGE interneuron migration deficit in *SLC6A1* mutant MGEOs and assembloids. Although a migratory phenotype has not yet been reported in mouse models of *Slc6a1* DEE, a recent study of AAV9/h*SLC6A1* gene therapy administered at various timepoints from P0 to P28 found that only administration at P0 fully rescued abnormal behavioral and electrophysiological phenotypes^89^. These findings suggest that events occuring during early brain development underlie the pathogenic phenotypes in *SLC6A1* DEE and highlight the importance of early intervention to treat this disorder. Our results support these findings by identifying that full and prolonged expression of GAT1 throughout the migratory period is required for the proper migration of MGE interneurons.

Interestingly, we could not replicate migration defects in WT MGEOs using a highly specific GAT1 blocker, NO711, applied at a concentration established to fully block GABA transport^90–92^. This finding suggests that early migration defects do not involve GAT1 transporter dysfunction. Indeed, prior work suggests that GAT1 may not function as a transporter until later developmental timepoints^71,93^, and one study showed that blockade of GAT1 in E14.5 mouse telencephalic slice co-cultures did not alter the pattern of MGE tangentially migrating cells^78^. While the exact mechanism of how GAT1 regulates early interneuron migration requires further investigation, studies in the field of oncology have also identified GAT1 as a modulator of cancer cell migration and invasion^94,95^. Knockdown of GAT1 in breast cancer cells reduces their migration and metastasis, potentially acting through the PI3K/AKT pathway^94^. Irrespective of mechanism, the migratory deficits of MGE cells that we identified likely contribute to seizures and ID associated with *SLC6A1* DEE. Because human MGE interneurons continue to migrate and mature until approximately 2-3 years of age^96–98^, they are a plausible target for early treatment interventions. Further work is needed to identify GAT1 interacting partners and potential druggable targets to rescue migration.

Our novel hPSC-derived MGEO model provides a valuable tool to study MGE-derived cortical interneuron development and for modeling of interneuron-related phenotypes in genetic epilepsies and related neurodevelopmental disorders (see **Figure S7** for expression of genes in MGEOs that are commonly implicated in neurodevelopmental disorders). In particular, our protocol offers the unique opportunity to study the intrinsic developmental programs of human PV cortical interneurons and their associated disorders. In addition to disease modeling, MGEOs may also serve as a potential source of MGE interneuron progenitors for therapeutic grafting to treat disorders associated with MGE-derived interneuron dysfunction, including epilepsies, ASD and neurodegenerative disorders.

### Limitations of this study

Some cell line variability appeared in the MGEO electrophysiological characterizations, particularly in their maturation timelines, with the fibroblast-derived line (2E) displaying slightly delayed electrophysiological development compared to the embryonic (H9) and blood-derived (79B) lines. These differences highlight the importance of validating hPSC lines and the need to utilize proper isogenic or cell-line-origin matched controls when modeling diseases. We are currently working on methods to further shorten the maturation period needed for PV expression. Additionally, MGEOs must be individually removed from the thick extracellular matrix manually at 10-12 DIV, limiting large scalability of this method. Further protocol optimizations are therefore needed to identify an efficient way to overcome this rate-limiting protocol step. Lastly, our MEA recordings were performed at a sampling rate of 12.5 kHz, limiting the temporal resolution needed to accurately classify fast spike waveform features (e.g. trough to peak) faithfully. To increase the specificity of cell-type identification in future experiments, incorporating a reporter line that specifically labels PV cortical interneurons would enable selective targeting of these cells. Regardless of these limitations, our findings demonstrate MGEO-CO assembloid network maturation, including increased firing rates, burst dynamics, and the presence of functionally integrated MGEO-derived inhibitory neurons.

## Resource availability

### Lead contact

Further information and requests for resources and reagents should be directed to the lead contact, Jack Parent.

### Materials availability

Materials used in this study are available upon request to the lead contact.

### Data and code availability

- The data generated during this study are available from the lead contact upon request. Raw single cell RNAseq data will be deposited to GEO upon publication.
- The code used for scRNAseq analysis and spike sorting analysis will be available upon request.
- Any additional information required to reanalyze the data reported in this paper is available from the lead contact upon request.

## Acknowledgments

Research reported in this publication was supported by the University of Michigan Advanced Genomics Core, the UM Single Cell Spatial Analysis Program and the National Cancer Institute under Award Number P30CA046592 by the use of the following Cancer Center Shared Resource: Single Cell and Spatial Analysis Shared Resource. We would like to thank the University of Michigan Viral Vector Core for generating the viral reagents. We also gratefully acknowledge Paul Jenkins for the generous gift of the AnkG antibody, and thank Maria Faraj for assisting with the AMD3100 migration experiments. This work was also supported by the National Institutes of Health (NIH; NS117170-JMP, NS129850-JF, NS076752-LLI), the National Science Foundation (NSF; EFMA 2422149-JF), the Simons Foundation Autism Research Initiative (JMP), the NSF Graduate Research Fellowship (DGE-2241144-MCV), the NIH Diversity Specialized Predoctoral to Postdoctoral Advancement in Neuroscience (D-SPAN) F99/K00 transition award (1F99NS135817-MCV), University of Michigan Rackham Merit Fellowship (MCV), T32 University of Michigan Training Program in Translational Research NIH-NIGMS Pre-doctoral Fellowship (5T32GM141840-02, 1T32GM141840-01-MPW), and the NIH Early Stage Training in the Neurosciences Training Grant (5T32NS076401-20-MPW). Some illustrations were created using BioRender.

## Author contributions

Conceptualization: M.C.V., M.P.W., J.M.P.; Methodology: M.C.V., M.P.W., J.M.P.; Data generation and analysis: M.C.V., M.P.W., T.T., L.G.; scRNA-seq data processing and analysis: J.B., M.C.V., M.P.W.; Electrophysiology acquisition and analysis: Y.Y., M.C.V., E.L.C.; Disease model experimental design: M.C.V., A.M.T., M.U., J.M.P; Writing of original draft: M.C.V, M.P.W with contributions from E.L.C., Y.Y., J.B.; Editing of original draft: J.M.P., M.U., A.M.T.; Review of final draft: all authors; Contribution of resources: J.M.P., M.U., L.L.I

## Declaration of interests

The authors have no financial or other competing interests to declare.

## STAR⍰Methods

### Key resources table

| REAGENT or RESOURCE | SOURCE | IDENTIFIER |
| --- | --- | --- |
| **Antibodies** |  |  |
| Mouse Monoclonal Anti-**LHX6** Antibody | Santa Cruz Biotechnology | Cat# sc-271433; RRID: AB_10649856 |
| Rabbit Polyclonal Anti- **Calretinin** Antibody | Millipore | Cat# AB5054; RRID: AB_2068506 |
| Rat Monoclonal Anti-**Somatostatin** Antibody | Millipore | Cat# MAB354; RRID: AB_2255365 |
| Mouse Monoclonal Anti-**Somatostatin** Antibody | Santa Cruz Biotechnology | Cat# sc-55565; RRID: AB_831726 |
| Rabbit Polyclonal anti-**Glial Fibrillary Acidic Protein** (GFAP) Antibody | Agilent | Cat# Z0334; RRID: AB_10013382 |
| Rabbit Polyclonal anti-**OLIG2** Antibody | Millipore | Cat# AB9610; RRID: AB_570666 |
| Rat Monoclonal anti-**Myelin Basic Protein** (MBP), aa 82-87 Antibody | Millipore | Cat# MAB386; RRID: AB_94975 |
| Rabbit Polyclonal Anti-**Parvalbumin** Antibody | Swant | Cat# PV27; RRID: AB_2631173 |
| Rabbit Polyclonal anti-**K<sub>v</sub>3.1b** (KCNC1) Antibody | Alamone | Cat# APC-014; RRID: AB_2040166 |
| Mouse Monoclonal Anti-**MEF2C** Antibody | Novus Biologicals | Cat# NBP2-00493; RRID: AB_3245039 |
| Wisteria floribunda lectin (**WFA**, WFL), Biotinylated | Vector Laboratories | Cat# B-1355; RRID: AB_2336874 |
| Goat anti-**Ankyrin G** Antibody | Gift from Paul Jenkins Lab | N/A |
| Rabbit Polyclonal Anti-**vGAT** Antibody | Synaptic Systems | Cat# 131 003; RRID: AB_887869 |
| Mouse Monoclonal Anti-**Map2ab** Antibody | Sigma-Aldrich | Cat# M4403; RRID: AB_477193 |
| Mouse Monoclonal Anti-**Gephyrin** Antibody | Synaptic Systems | Cat# 147 011; RRID: AB_887717 |
| Anti-Mouse Polyclonal Anti-**Pax6** Antibody | MBL International | Cat# PD022; RRID: AB_1520876 |
| Mouse Monoclonal Anti-**SHH** Antibody | Santa Cruz Biotechnology | Cat# sc-365112; RRID: AB_10709580 |
| Guinea Pig Polyclonal Anti-**Rx** Antibody | Takara | Cat# M229;<br>RRID: AB_2783559 |
| Mouse Monoclonal Anti-**NKX2.1** Antibody | Millipore | Cat# MAB5460;<br>RRID: AB_571072 |
| Mouse Monoclonal Anti-**Tuj1** Antibody | Covance | Cat# MMS-435p;<br>RRID: AB_2313773 |
| Rabbit Polyclonal Anti-**Doublecortin** Antibody | Abcam | Cat# AB18723;<br>RRID: AB_732011 |
| Goat anti-Mouse IgG (H+L) Highly Cross-Adsorbed Secondary Antibody, Alexa Fluor 488 | Thermo Fisher Scientific | Cat# A-11029;<br>RRID: AB_2534088 |
| Goat anti-Mouse IgG (H+L) Cross-Adsorbed Secondary Antibody, Alexa Fluor 568 | Thermo Fisher Scientific | Cat# A-11004;<br>RRID: AB_2534072 |
| Goat anti-Mouse IgG (H+L) Highly Cross-Adsorbed Secondary Antibody, Alexa Fluor 647 | Thermo Fisher Scientific | Cat# A-21236;<br>RRID: AB_2535805 |
| Goat anti-Mouse IgG (H+L) Cross-Adsorbed Secondary Antibody, Alexa Fluor 405 | Thermo Fisher Scientific | Cat# A-31553;<br>RRID: AB_221604 |
| Goat anti-Rabbit IgG (H+L) Cross-Adsorbed Secondary Antibody, Alexa Fluor 488 | Thermo Fisher Scientific | Cat# A-11008;<br>RRID: AB_143165 |
| Goat anti-Rabbit IgG (H+L) Cross-Adsorbed Secondary Antibody, Alexa Fluor 568 | Thermo Fisher Scientific | Cat# A-11011;<br>RRID: AB_143157 |
| Goat anti-Rabbit IgG (H+L) Highly Cross-Adsorbed Secondary Antibody, Alexa Fluor 647 | Thermo Fisher Scientific | Cat# A-21245;<br>RRID: AB_2535813 |
| Goat anti-Rabbit IgG (H+L) Cross-Adsorbed Secondary Antibody, Alexa Fluor 405 | Thermo Fisher Scientific | Cat# A-31556;<br>RRID: AB_221605 |
| Goat anti-Guinea Pig IgG (H+L) Highly Cross-Adsorbed Secondary Antibody, Alexa Fluor 647 | Thermo Fisher Scientific | Cat# A-21450;<br>RRID: AB_2535867 |
| Donkey anti-Goat IgG (H+L) Cross-Adsorbed Secondary Antibody, Alexa Fluor 488 | Thermo Fisher Scientific | Cat# A-11055;<br>RRID: AB_2534102 |
| Donkey anti-Rabbit IgG (H+L) Highly Cross-Adsorbed Secondary Antibody, Alexa Fluor 488 | Thermo Fisher Scientific | Cat# A-21206;<br>RRID: AB_2535792 |
| **Bacterial and virus strains** |  |  |
| pAAV_BiPVe3_dTomato_nls-dTomato | Furlanis et al.<sup>99</sup> | Addgene 213940 |
| pAAV-mDLX-GFP_Fishell-1 | Dimidschstein et al.<sup>100</sup> | Addgene 83900-AAV1 |
| pAAV-CAG-tdTomato (codon diversified) | Edward Boyden | Addgene 59462-AAV1 |
| **Chemicals, peptides, and recombinant proteins** |  |  |
| DMEM/F12 | Gibco | Cat# 11330032 |
| mTeSR1 | STEMCELL Tech | Cat# 85850 |
| mTeSR Plus | STEMCELL Tech | Cat# 100-0276 |
| Geltrex | Thermo Fisher Scientific | Cat# A1413302 |
| AccuGENE 0.5M EDTA Solution | Lonza | Cat# 51201 |
| Rho-kinase inhibitor (Y-27632) | Tocris | Cat# 1254 |
| StemPro™ Accutase™ | Thermo Fisher Scientific | Cat# A1110501 |
| Neurobasal | Gibco | Cat# 21103049 |
| N-2 Supplement | Gibco | Cat# 17502048 |
| B-27 Supplement | Gibco | Cat# 17504044 |
| B-27 Supplement minus Vitamin A | Gibco | Cat# 12587010 |
| DMH1 | Tocris | Cat# 4126 |
| XAV939 | Cayman Chemical | Cat# 13596 |
| SB-431542 | Cayman Chemical | Cat# 13031 |
| Cyclopamine | Cayman Chemical | Cat# 11321 |
| L7 Passaging Solution | Lonza | Cat# FP-5013 |
| CHIR99021 | Cayman Chemical | Cat# 13122 |
| SAG | Selleckchem | Cat# S7779 |
| Human FGF-basic (FGF-2/bFGF) (aa 1-155)<br>Recombinant Protein | Gibco | Cat# PHG0261 |
| Human Albumin | Sigma | Cat# A7223-1G |
| Human Brain Derived Neurotrophic Factor<br>(BDNF) | Peprotech | Cat# 450-02-1MG |
| Human Neurotrophic factor-3 (NT3) | Peprotech | Cat# 450-03-50UG |
| MEM Non-Essential Amino Acids (MEM NEAA) | Gibco | Cat# 1140-050 |
| Penicillin Streptomycin | Gibco | Cat# 14140-122 |
| Glutamax (100X) | Gibco | Cat# 35050-061 |
| BrainPhys Neuronal Medium with SM1 Kit | STEMCELL Tech | Cat# 5792 |
| L-Ascorbic acid | Sigma | Cat# A92902-500G |
| Human GDNF | Peprotech | Cat# 450-10-1MG |
| HEPES (1M) Buffer Solution | Gibco | Cat# 15630-080 |
| AMD3100 | Tocris | Cat# 3299 |
| SC-79 | Cayman Chemical | Cat# 14972 |
| NNC-711 (NO711) | Abcam | Cat# ab120364 |
| Streptavidin, Alexa Fluor™ 647 Conjugate | Thermo Fisher Scientific | Cat# S21374 |
| **Critical commercial assays** |  |  |
| RNeasy mini kit | Qiagen | Cat# 74104 |
| RNase-Free DNase Set | Qiagen | Cat# 79254 |
| SuperScript™ III First-Strand Synthesis<br>SuperMix kit | Invitrogen | Cat# 11752-050 |
| Power SYBR Green PCR Master Mix | Thermo Fisher Scientific | Cat# 4367659 |
| Chromium Next GEM Single Cell Fixed RNA<br>Sample Preparation Kit | 10x Genomics | Cat# 1000414 |

| Deposited data |  |  |
| --- | --- | --- |
| Raw MGEO scRNAseq data | This paper | Available upon publication |
| Human reference genome NCBI build 38, patch 13 (GRCh38.p13) | Genome Reference Consortium | http://www.ncbi.nlm.nih.gov/projects/genome/assembly/grc/human/ |
| scRNAseq from human ganglionic eminences at GW09, GW11, GW12, GW13, GW16, & GW18 | Shi et al.<sup>11</sup> | GEO: GSE135827 |
| scRNA-seq for hMGEO organoids derived from hPSCs | Xiang et al.<sup>101</sup> | GEO: GSE97882 |
| scRNA-seq for hSS organoids derived from human iPSCs | Birey et al.<sup>102</sup> | GEO: GSE93811 and GSE96045 |
| Experimental models: Cell lines |  |  |
| Con2E (2E) Male Fibroblast-derived iPSC line | Tidball, et al.<sup>103,104</sup> | N/A |
| 79B Female PBMC-derived iPSC line | Mojica-Perez, et al.<sup>105</sup> | N/A |
| H9 Female Embryonic hESC line | Thomas et al.<sup>106</sup> | WiCell cat# WA09<br>RRID:CVCL_9773 |
| Oligonucleotides |  |  |
| See Table S1 for RT-qPCR primer sequences |  |  |
| Software and algorithms |  |  |
| BioRender | N/A | RRID:SCR_018361 |
| Fiji | Schindelin et al.<sup>107</sup> | RRID:SCR_002285 |
| RStudio (v4.4.1) | Posit team | RRID:SCR_000432 |
| Seurat (v4.2.3) | Hao*, Hao*, et al.<sup>108</sup> | RRID:SCR_000432 |
| Cell Ranger Single-Cell software suite (v7.1.0) | 10x Genomics | RRID:SCR_017344 |
| Loupe Browser | 10x Genomics | RRID:SCR_018555 |
| GraphPad Prism 10 | N/A | RRID:SCR_002798 |
| BCL Convert Conversion Software (bcl2fastq) v4.0 | Illumina | RRID:SCR_015058 |
| Other |  |  |
| StemPro EZ passage tool | Thermo Fisher Scientific | Cat# 23181010 |
| STRIPPER® Micropipetter | Cooper Surgical | Cat# MXL3-STR |
| 275µM tips for STRIPPER Micropipetter | Cooper Surgical | Cat# MXL3-275 |
| O.C.T. | Fisher Healthcare | Cat# 4585 |
| Tissue-Tek Cryomolds | Sakura | Cat# 32663 |
| Costar Ultra-low attachment 96-well round bottom plates | Corning | Cat# CLS7007-24EA |
| Costar 24-well Ultra-Low attachment plates | Corning | Cat# 3473 |

### Human Pluripotent Stem Cell Culture

All hPSC experiments were conducted with prior approval from the University of Michigan Human Pluripotent Stem Cell Research Oversight (HPSCRO) Committee. To ensure the reproducibility of our MGEO protocol, experiments were replicated using three hPSC lines, one hESC line and two iPSC lines one each derived from fibroblasts and peripheral blood mononuclear cells, and including both male and female lines.

Cultures were maintained in incubators dedicated to hPSCs, which were kept at 37°C with humidity control and 5% CO2. Prior to organoid differentiation, stem cells were maintained in mTeSR-1 (STEMCELL Technologies, Cat# 85850) or mTeSR-Plus (STEMCELL Technologies, Cat # 100-0276) feeder-free medium on 6-well tissue culture plates coated with 1:100 Geltrex (Thermo Fisher scientific A1413302). Cells were regularly passaged when they reached approximately 60-70% confluency using 0.8mM EDTA. Cell lines were routinely tested for mycoplasma and monitored for morphology and spontaneous differentiation. The genomic integrity and pluripotency of these cell lines was confirmed by Single Nucleotide Polymorphism (SNP) DNA Microarray and immunostaining for pluripotency markers, respectively^38,39,113,114^.

### Generation of SLC6A1 DEE lines

To generate *SLC6A1*gene edited iPSC lines Human foreskin fibroblasts were reprogrammed into iPSCs with concurrent CRISPR/Cas9 gene editing^77^ targeted to generate out-of-frame indels in exon 7 of the *SLC6A1* gene. Genotypes of heterozygous (Het), knock-out (compound Het) and isogenic control lines were confirmed by Sanger and next-generation sequencing, with SNP-ChIP analysis assessing chromosomal integrity (data not shown). To generate the SLC6A1 rescue line, the PiggyBac transposase system was used to stably integrate vectors containing doxycycline-inducible, codon optimized, WT human *SLC6A1* into a knockout *SLC6A1* iPSC line (KOi). Increase in GAT1 mRNA was confirmed in KOi MGEOs using RT-qPCR and conditions using the KOi line without the addition of dox was used as a negative control. To generate patient derived iPSC lines, blood samples were acquired with consent from *SLC6A1* patients within the University of Michigan Health system. An Erythroid Progenitor Reprogramming kit (Stem cell technologies, 05924) was used to isolate and expand erythroid progenitors from peripheral blood mononuclear cells and transfect a reprogramming vector cocktail containing: pCXLE-hSK (SOX2, KLF4); pCXLE-hUL (L-Myc); pCXL-hOCT3/4 (Oct3/4). Pluripotency of all iPSC lines were validated using RT-qPCR and immunostaining of pluripotency markers.

### MGEO generation

Self-organizing single rosette MGEOs were generated from a hPSC monolayer by modifying our SOSR-CO protocol (Tidball et al., 2023b). For the initial formation of the monolayer, iPSC lines were dissociated using Accutase (Innovative cell) and plated onto Geltrex-coated (1:50 dilution in DMEM/F12) 12-well plates at 3–4 × 10^5^ cells/mL in mTeSR1 with 10 μM Rho-kinase inhibitor (Y-27632; Tocris, 1254). Media without the inhibitor was replaced daily until the cells reached 80%–100% confluency. The medium was then changed to 3N (50:50 DMEM/F12:Neurobasal with N2 and B27 supplements) without vitamin A with 2 μM DMH1 (Tocris, 4126), 2 μM XAV939 (Cayman Chemical, 13596), and 10 μM SB431542 (Cayman Chemical, 13031) (2 mL of media per well). On DIV 4, media was removed, and the monolayer was incubated for 1 min with L7 hPSC passaging solution (Lonza). After aspirating the L7 solution, the monolayer was cut into squares using a StemPro EZ passage tool. The squares were sprayed off the bottom of the culture plate with 1 mL of pre-prepared culture media with a P1000 micropipette.

Approximately 200 μL of resuspended squares were then transferred into each well of a 96-well plate preincubated at 37°C for 30 min with 40 μL of 100% Geltrex solution/well. Each monolayer will seed up to 16 wells containing approximately 20-40 organoids per well. Additional media was added to each well the day after plating without media change. Half-media changes were performed daily beginning 48 h post plating. On day 6 (2 days after cutting), human FGF2 (20 ng/ml, Gibco, PHG0261) containing 0.1% human serum albumin was added to the culture media to support cell survival. On day 10, a 100% media change was performed with 3N media plus 100 nM SAG (Selleckchem, S7779) and FGF2, without DMH1, XAV939 and SB431542. Between days 10 and 12 (based upon size), the organoids were individually removed using a STRIPPER Micropipetter (Cooper Surgical) with 275 μm tips and placed in suspension individually in the wells of a low-adherence U-bottom 96-well plate with 200 μL of 3N medium without vitamin A. The ideal diameter for picking the MGEOs is ∼250 μM. Larger sizes may produce additional rosettes, and smaller sizes tend to have suboptimal survival. Starting on day 12, half-media changes (100 µL/well) of 3N with vitamin A, 100 nM SAG, BDNF (20 ng/mL), and NT3 (20 ng/mL) without FGF2 were repeated every other day until day 18 when SAG was removed from the culture medium. After 35 days of differentiation, 3N media with vitamin A (but without BDNF and NT3) was used for half-media changes every other day. At this time, MGEOs were transferred to low-adherence 24-well plates for a necessary increase in media volume due to expanded organoid size. Beginning day 90, MGEOs were cultured in BrainPhys media (Stemcell technologies, 05790) containing N2 supplement, NeuroCult™ SM1 Neuronal Supplement (Stemcell technologies, 05711), BDNF (20 ng/mL), GDNF (20 ng/mL), and Ascorbic acid (200nM). Self-organizing single rosette cortical organoids (SOSR-COs) were generated as described^38^.

### Cryopreservation, sectioning and immunocytochemistry

Organoids were collected, washed with phosphate-buffered saline (PBS), and incubated in 4% paraformaldehyde (PFA) diluted in PBS at 4°C for 30 min (for 1-month organoids) or an hour (for older organoids). After three 10-min PBS washes, organoids were incubated overnight at 4°C in a 30% sucrose (in PBS) solution, embedded in cryomolds (Tissue-Tek 4566) using O.C.T compound (Fisher Healthcare 4585), frozen on dry ice, and stored at-80°C. Frozen organoid blocks were sectioned onto Superfrost Plus glass slides (ThermoFisher 12-550-15) at a thickness of 20 μM using a Leica CM1850 cryostat. After drying at room temperature (RT) for at least 2 h, slides were stored at - 20°C until use. For immunostaining, organoid sections were washed three times with PBS, permeabilized for 20 min in PBS with 0.2% Triton-X100 at RT, and incubated in blocking buffer (PBS with 0.05% Tween-20, 5% normal goat serum, and 1% bovine serum albumin (BSA)) for 1 h at RT. Normal horse serum was used in blocking buffer for immunostaining using goat primary antibodies. Fixed and permeabilized sections were incubated overnight at 4°C in a humidified chamber with primary antibodies diluted in blocking buffer. Sections were then washed 3 times in PBST (PBS with 0.05% Tween-20) and incubated for 2 h at RT with Alexa-Flour secondary antibodies diluted 1:500 in blocking buffer. Sections were washed once with PBST for 10 min and incubated with bisbenzimide for 5 min to label nuclei. After 3 x 10 min washes of PBST, coverslips were mounted onto slides using Glycergel mounting medium (Agilent Dako C0563). Once set, the edges of the slides were sealed using clear nail polish. Slides containing immunolabeled organoid sections were imaged using an Andor BC43 benchtop spinning disc confocal microscope (Oxford Instruments). Acquired images were analyzed in Fiji.

### RT-qPCR

For reverse transcriptase quantitative PCR (RT-qPCR), total RNA was extracted using the miRNeasy Mini Kit (Qiagen 217004) as per manufacturer’s protocols (with optional DNase1 digestion step), with at least four organoids pooled per sample. Template RNA (100 ng) was used to synthesize cDNA using the SuperScript™ III First-Strand Synthesis SuperMix kit (Invitrogen 11752-050). RT-qPCR was performed as per manufacturer protocols, using 2 ng of cDNA and Power SYBR Green PCR Master Mix (Thermofisher 4367659) with an AppliedBiosystems QuantStudio3 thermocycler (see **Supplemental Table S1** for primer sequences). Fold changes were calculated using the 2^-ΔΔCt^ method, normalizing to the average undifferentiated 2E iPSC levels from n=3-4 experiments. Samples were run in triplicate and GAPDH was used as an internal control. Undetectable levels of mRNA were set to a CT value of 40 for analysis, which was performed in Microsoft Excel and graphed using GraphPad Prism.

### Fixation of whole MGE organoids

Whole organoid fixation for downstream scRNA-seq analysis of MGEOs was conducted using the 10X Genomics Chromium Next GEM Single Cell Fixed RNA Sample Preparation Kit (10X Genomics, 1000414). All reagents described below were supplied with the kit and the protocol was carried out as described on the 10X Genomics website. In brief, 4-8 organoids (depending on age/size) were collected using a P1000 micropipette with a cut pipette tip and transferred to a 1.5 mL Eppendorf tube.

Organoids were allowed to sink to the bottom of the tube and the supernatant media was carefully aspirated from the tube without disturbing the organoids. Organoids were rinsed once with 1 ml of PBS, the PBS was removed from the tube and 500 ml of Fixation Buffer was added and incubated for 24 h at 4°C. The next day, the Fixation Buffer was removed without disturbing the organoids and the organoids were washed with 1 mL of chilled PBS. PBS was removed and 500 µL of Quenching Buffer was added to the tube containing the fixed organoids. Enhancer (50 µL) was added and pipette mixed. Glycerol (137.5 µL of a 50% solution) was then added and pipette mixed for a final concertation of 10% glycerol. Samples were stored in-80°C until they were sent to the University of Michigan’s Advanced Genomics core for further processing and analysis.

### MGEO tissue dissociation, library prep, and sequencing

Tissue dissociation, library prep and next-generation sequencing was carried out in the Advanced Genomics Core at the University of Michigan. Fixed organoid tissue samples were dissociated, then individual samples were barcoded and pooled together for sequencing. Cells were processed through the 10x Genomics Chromium system (10x Genomics). 10x Genomics v.3 libraries were prepared according to the manufacturer’s instructions. Libraries were then sequenced using paired-end sequencing with a mean coverage ranging from 15,000 to 39,000 raw reads per cell using an Illumina NovaSeqXPlus. Day 90 pooled samples were subjected to 28×90bp of sequencing according to the manufacturer’s protocol (Illumina NovaSeqXPlus). scRNA-seq data were aligned and quantified using Cell Ranger Single-Cell software suite (v.7.1.0, 10x Genomics) against Homo sapiens (human) genome assembly GRCh38.p13 from ENSEMBL. BCL Convert Conversion Software v4.0 (Illumina) was used to generate de-multiplexed Fastq files.

### Analysis of scRNAseq data

Analysis of scRNA-seq data was done as previously described^115^. Briefly, the data were analyzed using the R package Seurat (v.4.2.3, https://satijalab.org/seurat/)). Datasets were filtered, normalized, and scaled. Differences between G2/M phase and S phase scores were regressed out during the scaling process. The dataset further went through PCA analysis and was embedded into low-dimensional space using UMAP algorithm for visualization. Clusters were identified using shared nearest neighbor modularity optimization-based clustering algorithm. The clusters were annotated based on expression of known marker genes and genes found to identify specific cell types in human fetal MGE data sets ^11,15,28^.

To compare our day 90 MGEO scRNA-seq data to human fetal GE expression, we used the Shi *et al.* dataset ^11^, which includes scRNA-seq data from MGE, LGE, and CGE human fetal tissue at GW 9, 12, 16, and 18. A PCA plot was used to determine how closely related the gene expression profile of our MGEO samples were to the GW9-19 human dataset’s regional and cell type clusters, and to determine the GW age to which the MGEOs were most similar. A Pearson correlation was used to compare day 90 MGEO neuronal clusters 1-7 to human fetal MGE subclusters. To assess regional specificity and determine the extent to which MGEOs more closely recapitulate MGE-like expression markers, we compared the percentage of cells expressing key regional and interneuron gene markers between our MGEO scRNA-seq dataset, established subpallial organoid protocols^32,33^, and the Shi *et al*. human MGE cluster.

The number and percentage of cells expressing key regional and interneuron gene markers in each dataset was calculated and plotted using a custom R function. In brief, gene expression percentages were calculated across different Seurat objects, combined into a data frame, transformed using the melt() function, and visualized for comparison across datasets using ggplot2. To visualize the number of neurons expressing one or more interneuron subtype-specific markers, neuronal clusters (clusters 1-7) were extracted from the day 90 MGEO data set and the top five subtype specific gene markers (based on the percentage of neurons expressing each marker) were selected for multi gene expression analysis. All valid gene combinations were identified using a custom R function and the number of neurons with single or multi gene expression profiles was plotted using ggplot2.

### Viral labeling and generation of cortical-MGEO assembloids

To visualize migrating neurons in assembloids, cortical organoids (SOSR-COs) were labeled with codon diversified AAV1-CAG-tdTomato (Addgene # 59462), while interneuron progenitors and interneurons were labeled in MGEOs using AAV1-mDlx-GFP virus (Addgene # 83900) prior to fusion. Briefly, 46 DIV organoids were transferred into 1.5 mL Eppendorf tubes and incubated overnight (37°C, 5% CO2) in 250 μL of 3N with vitamin A organoid culture media containing diluted virus (1:50). Fresh 3N media with Vitamin A (750 μL) was added the next day. The following day, organoids were washed with PBS to remove residual virus, then transferred to ultra-low attachment plates with fresh culture media for maintenance until fusion into assembloids approximately one week later. To generate MGEO-cortical assembloids, SOSR-CO and MGEO regions with the brightest viral reporter expression were placed in contact with each other along the bottom edge of a 48-or 24-well plate. Plates were tilted at an angle to ensure the organoids maintained contact with each other, and left undisturbed for at least 24 hours. At later stages, PV basket cells were labeled in assembloids with AAV1-BiPVe3-dTomato_nls-dTomato virus (Addgene # 213940) using the same methods as above.

### 2D assay of MGEO neuronal migration

To interrogate the migration of interneurons from 3D MGEOs onto 2D extra-cellular matrix, 21 DIV organoids were plated into individual wells of a 1:50 Geltrex-coated, thin-bottom, black-walled 96-well culture plate (PerkinElmer 6055300) containing 3N+A media. Organoids were gently moved to the center of each well using a pipette tip and allowed to settle to the bottom of the well. Automated live imaging of migration in phase or brightfield was performed using an Incucyte S3 Live Cell Analysis System or Andor BC43 benchtop spinning disc confocal microscope. Using the Incucyte, multiple 10x images were acquired per well at 15-min intervals over 48 h, starting immediately after plating. Andor confocal microscope images were collected at 5-10 min intervals over 15 h, beginning at least 2 h post plating. For both imaging methods, cultures were maintained at 37°C and 5% CO2. Plates were fixed with 4% PFA at 48 h post-plating for immunostaining, which was performed as described above.

To block CXCR4, a receptor that regulates the migration of interneurons in the developing neocortex^66,67^, the CXCR4 antagonist AMD3100 (100 nM) was added to organoid media^32^. For the “chronic” exposure condition, AMD3100 was added beginning 6 days prior to the migration assay (24 DIV), with a half media change containing fresh 100 nM drug performed every 2-3 d. For the “acute” groups, 100 nM of drug was added at the time of plating for the migration assay (30 DIV). Incucyte images were taken as described above and Fiji ImageJ was used to quantify the area of cell migration (onto ECM) between control, acute, and chronic conditions at 48-h post plating. The organoid area and spots of debris were excluded prior to quantification. Experiments were performed using 8 MGEOs per condition from 2 differentiations derived from the 79B hPSC line. Graphpad Prism was used to graph the migrated neuron area of each replicate organoid and for statistical analysis (see below). To block GAT1, the highly selective GAT1 antagonist NO711 (25 µM) was added to the organoid media of *SLC6A1* isogenic control MGEOs. Drug was refreshed daily with half media changes and was included for the entire duration of the migration assay from 21 DIV until 25 DIV. At 25 DIV, organoids and migrated cells were fixed with 4% PFA and immunostained with DCX and bisbenzimide to assess distance traveled of migrating interneurons.

Organoids were imaged on the Andor BC43 and analyzed in Fiji.

### Live imaging of assembloids, 3D migration, and quantification

To assess interneuron migration in a 3D environment, SOSR-COs and MGEOs were virally labeled and fused at 56 DIV as described above. Live imaging of GFP+, migrating cells was conducted using the Andor BC43 benchtop spinning disk confocal microscope. Migration in assembloids was imaged every 5 min for 15 h. Static confocal images of live assembloids were also collected at multiple timepoints between 24 h to 15 d post fusion to assess migration over time. mDLX-GFP labeled cells that migrated into the cortical organoid area were quantified in Fiji as the percent of GFP area in the CAG-TdTom area and plotted using Graphpad Prism.

Complementing our 2D neuronal migration experiments above, we exposed 79B-derived assembloids to the CXCR4 antagonist AMD3100 (or no drug control) 1 day post MGEO-CO fusion. We imaged and quantified the area of mDLX-GFP MGEO interneurons that migrated into CO organoid 24 h later, as described above.

### Multi-electrode array (MEA) recordings

Assembloids (at 20-34 wk differentiation) were plated individually in each well of a 6-well cytoview MEA plate (Axion Biosystems, Atlanta, GA, USA) coated with ploy-L-ornithine (PLO) (100 µg/mL) and Geltrex (1:100). As opposed to the conventional method of organoid MEA recording, wherein organoids grow chronically on the MEA plate, which alters their 3D cytoarchitecture over time, we transferred each organoid to the MEA plate immediately before recording and carefully removed the majority of the media after positioning the organoid on top of the electrodes. This method takes advantage of the adhesion properties of liquid, allowing the organoid to firmly contact the electrodes as the media is removed. After a 5-min recording, media was added back into the MEA plate and the organoid was gently removed from the MEA plate and maintained in suspension culture. The reproducibility of MEA recordings was verified by removing, replating, and then re-recording from the same organoid several times over a short time window and across multiple MEA wells to ensure consistency of signal (data not shown).

All recordings were performed using a Maestro MEA system and AxIS Software in Spontaneous Neural Configuration (Axion Biosystems). MEA signals were sampled at 12.5 kHz and bandpass filtered at 200-3000 Hz with Butterworth filtering to detect APs. APs were detected using a threshold of 5.5 times the standard deviation of the noise. Single electrode bursts were detected as a minimum of five APs occurring with a maximum interval of 100 ms. Network bursts were detected as a minimum of 50 APs with 100 ms maximum inter-AP interval observed on a minimum of 25% of active electrodes. Mean firing rate estimation was determined within a 10 s window and synchrony parameters were detected using a window size of 20 ms.

### Patch-clamp recording of organoid slices

Procedures for cutting organoid slices were similar to those used for acute mouse brain slices. The slicing solution contained (in mM): 110 sucrose; 62.5 NaCl; 2.5 KCl; 7.5 MgCl_2_; 1.25 KH2PO_4_; 26 NaHCO_3_; 0.5 CaCl_2_ and 20 D-glucose (pH 7.35-7.4 when saturated with 95% O2 /5% CO2 at room temperature of 22-25° C). To prepare fresh organoid slices, one assembloid (1-2 mm) was manually cut into 4-6 slices (∼300 µm) in ice-cold, oxygenated “slicing” solution saturated with 95% O_2_ /5% CO_2_. Slices were incubated in slicing solution for 15-20 min at RT and then in a mixture (1:1) of slicing solution and artificial cerebrospinal fluid (ACSF), the latter containing (in mM): 125 NaCl; 2.5 KCl; 1 MgCl_2_; 1.25 KH2PO_4_; 26 NaHCO_3_; 2 CaCl_2_ and 20 D-glucose, (pH 7.35-7.4), in a holding chamber aerated continuously with 95% O_2_ /5% CO_2_ at 35° C for 30 min before transfer to ACSF at RT.

Procedures for electrophysiological recordings in assembloid slices were similar to those used previously for acute mouse brain slices or for cultured iPSC-derived neurons^114,116–118^. An assembloid slice was transferred to a recording chamber where it was perfused (2-3 ml/min) with ACSF bubbled continuously with 95% O_2_ /5% CO_2_. The mDLX-GFP-labeled interneurons and CAG-tdTom-labeled pyramidal cells were visually selected using a NIKON E600FN upright microscope equipped with Nomarski optics (x 40 water immersion objective). For recording of AP firing, recording electrodes had resistances of 4-7 MΩ when filled with a potassium gluconate (K-gluconate)-based pipette solution consisting of (in mM): 140, K-gluconate; 4, NaCl; 0.5, CaCl_2_; 10, HEPES; 5, EGTA, 5 phophocreatine; 2, Mg-ATP and 0.4, GTP (pH 7.2-7.3 adjusted with KOH). Repetitive firing and AP frequencies from individual neurons were examined using the whole cell current-clamp recording technique. Repetitive AP firing was evoked by injection of a series of 1500 ms depolarizing currents varying from-60 pA to 200 pA at 10 pA-step from resting membrane potentials. Spontaneous AP firing was recorded in the gap-free model from resting membrane potentials. The pipette solution for spontaneous inhibitory synaptic current (sIPSC) recordings consisted of (in mM): 140, CsCl; 4, NaCl; 0.5, CaCl2; 10, HEPES; 5, EGTA, 5 phosphocreatine; 2, Mg-ATP and 0.4, GTP (pH 7.3 adjusted with CsOH). Signals were amplified with a Multiclamp 700B amplifier (Molecular Devices, Sunnyvale, CA), filtered at 2-4 kHz and digitized at 20 kHz for off-line analysis. Data were acquired with a Digidata 1440A interface and analyzed using pClamp11 offline. All experiments were carried out at RT (22-25°C).

### Statistics

GraphPad Prism 10.2.2 was used for all statistical analyses. In Figures 1 and 3, RT-qPCR data are presented as mean ± SEM, with each dot representing a single experimental replicate sample, which is composed of data from 4-8 pooled organoids. Migration data is graphed as mean ± SD with each dot being a datapoint from a single organoid. One-way ANOVA with Tukey multiple comparisons correction analysis was used to determine the statistical significance in migrated neuron area between control, acute, and chronic AMD3100 treatments at 48 h post plating for 2D migration assays. Student T-test statistical comparison was used to compare the effects of control vs AMD3100 treatment on the area of mDLX-GFP MGEO interneuron migration into the CO organoid area of assembloid at 48 h post-fusion. Experimental conditions were blinded during analysis. In all graphs, dot shapes denote the hPSC lines from which organoids were derived (circle for 79B, triangle for 2E, diamond for H9). For comparisons of MEA firing patterns (Figure 6), cumulative distribution of maximum firing rates and burst counts were compared via two-tailed Kolmogorov-Smirnov test. Firing rate per burst before and after bicuculline exposure was compared using two-tailed Mann-Whitney test.

---

**Supplementary Video S1: Rapid cell migration from a 21 DIV MGEO onto thin Geltrex in 2D migration assay.** Phase images were taken every 5 minutes (timestamps at bottom right, listed as hrs:mins:seconds) using an Incucyte S3 Live Cell Analysis System. Also see main figure 6.

**Supplemental Video S2: MGEO-derived interneurons show leading-edge branching migration patterns.** Time-lapse video of AAV-mDLX-GFP-labeled MGEO interneuron migration one day post fusion to a cortical organoid (CO) labeled with AAV-CAG-tdTom. Arrows track the soma of two neurons throughout the 4 hour time-course (time is listed as hr:min at top left). Organoids were derived from 2E male iPSC line and fused at 56 DIV. Also see **Figure 5**.

**Supplementary Video S3: MGEO-interneuron migration into cortical organoids 5-days post fusion in WT and PT assembloids.** Live confocal imaging of wildtype (left) and patient (right) assembloids fused at 56 DIV shows decreased migration of AAV1-mDLX-GFP-labeled MGEO interneurons (green) migrating into AAV-CAG-tdTom-labeled COs (red) in patient assembloids compared to wildtype. Imaged every 10 min for 15 h.

**Supplementary Figure S1:**
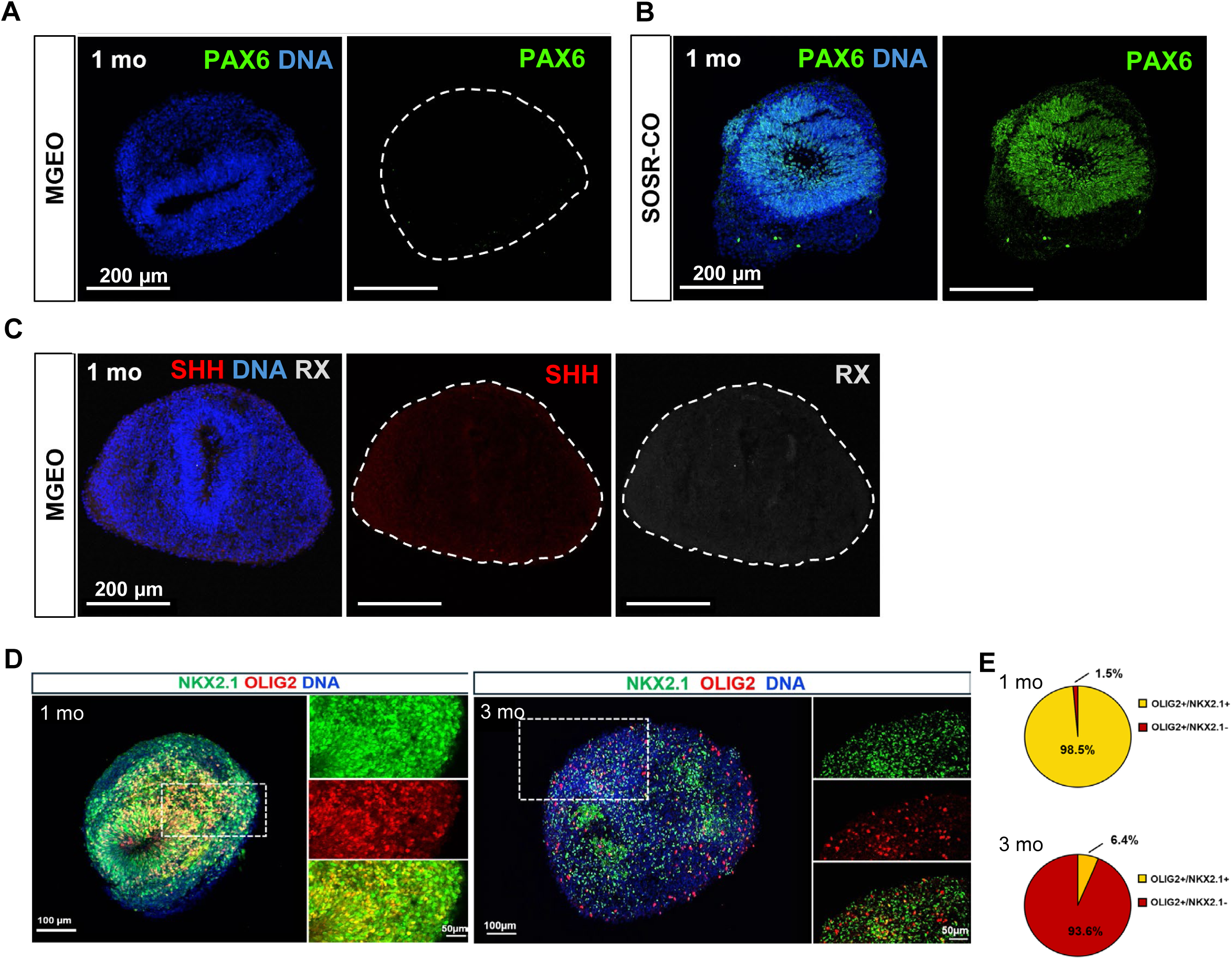
MGE regional specificity of MGEOs ((A, B) Immunostaining of PAX6 in 1-month (mo) MGEOs (A) and SOSR-COs (B). (C) 1-month MGEOs immunostained with SHH and RAX. (D) Immunostaining for NKX2.1 (green) and OLIG2 (red) at 1 month (left) and 3 months (right) shows co-expression at early timepoint, but not the latter. Boxed region indicates areas of magnification displayed on the right of each image. Bisbenzimide nuclear stain (DNA) is in blue for images in A-D. (E) Pie charts showing the percent of OLIG2 area that co-localizes with NKX2-1 (yellow) vs does not co-localize (red) in 1-and 3-month MGEOs.

**Supplemental Figure S2:**
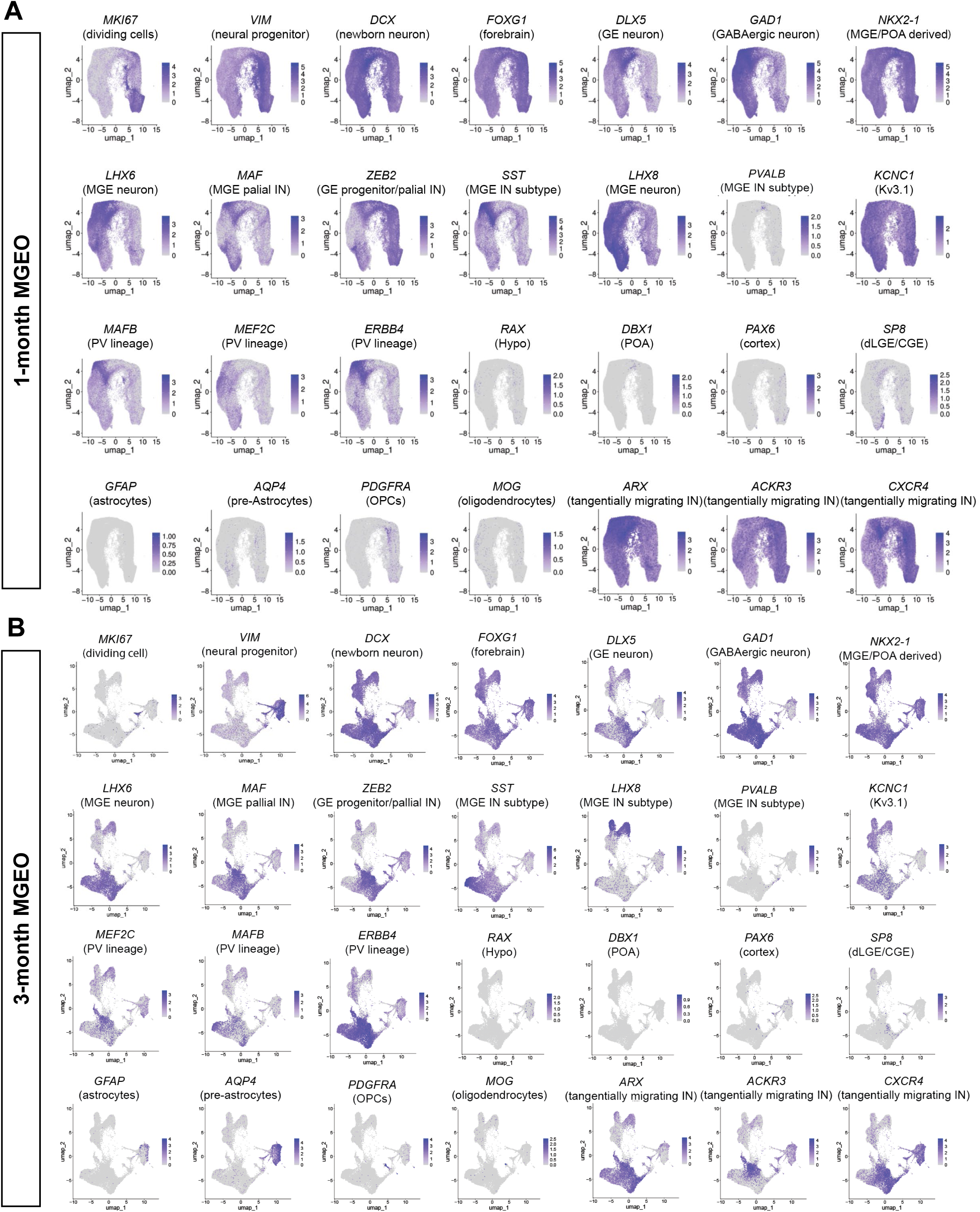
Expression of key markers in 1-month and 3-month MGEO scRNAseq data. Individual UMAP plots of selected genes expressed at 1 month (A) and 3 month (B). Color gradient denotes expression levels from none (light gray) to highest (dark purple) as depicted by the scale on the right of each plot.

**Supplementary Figure S3:**
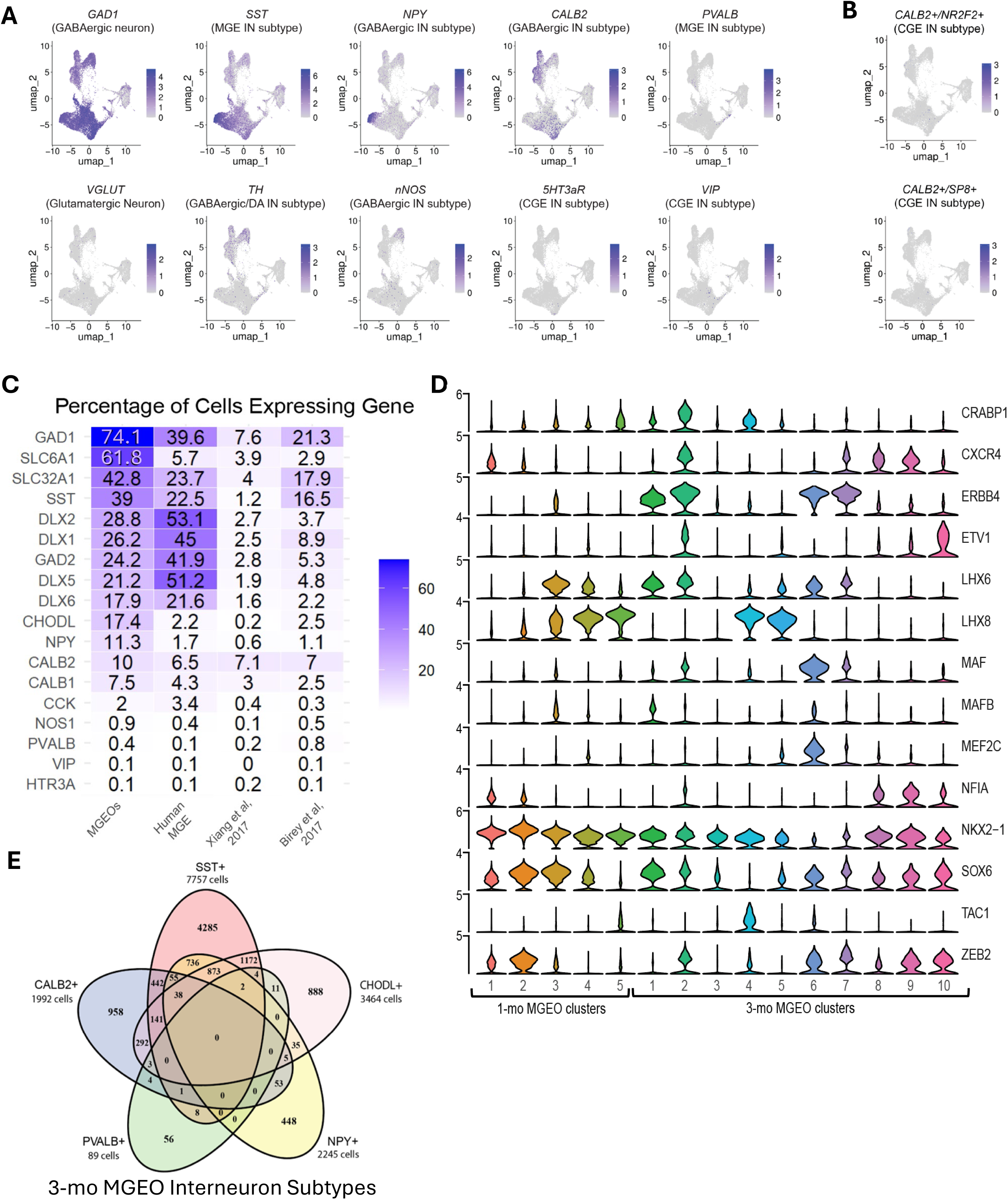
MGEO interneuron subtypes (A) UMAP feature plots showing expression of key GABAergic interneuron markers and lack of Glutamatergic and CGE cell types in 3-month MGEOs. (B) UMAP feature plots show minimal or no CALB2+ cells that co-express known CGE markers NR2F2 (top) or SP8 (bottom). (C) Heat map comparing the average percentage of cells expressing key GABAergic cell markers across scRNA-seq datasets from 3-month MGEOs, human fetal MGE (Shi et al., 2021; gestational weeks 9–18), and other subpallial organoids at similar timepoints (Xiang et al., 2017; Birey et al., 2017). (D) Stacked violin plots of MGE lineage markers grouped by MGEO age and cluster. (E) Venn diagram of scRNA-seq transcript co-expression between interneuron subtypes in 3-month MGEOs.

**Supplemental Figure S4:**
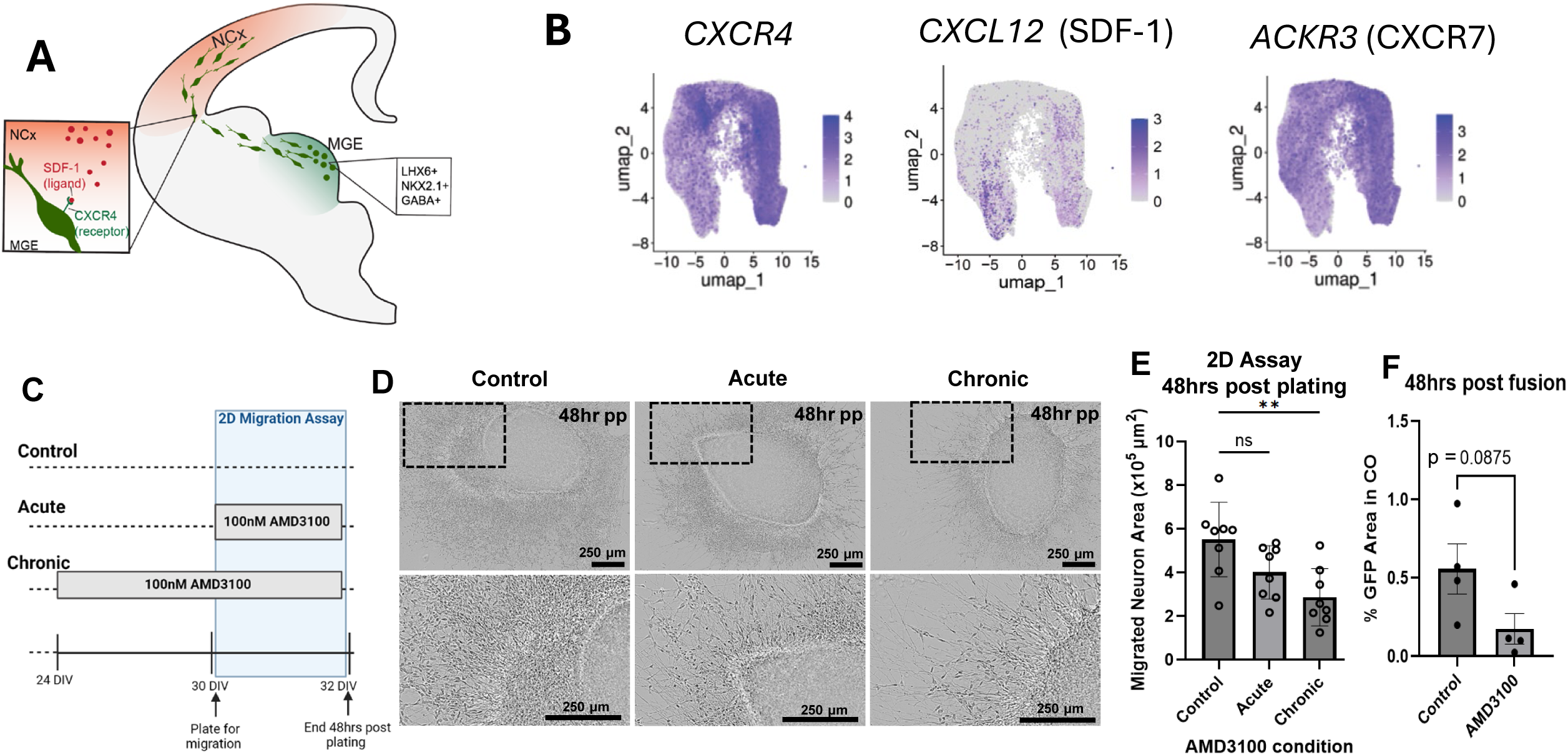
AMD3100 treatment reduces MGEO interneuron migration. (A) Schematic diagram showing SDF-1 binding to CXCR4 receptors to promote migration of MGE-derived GABAergic interneurons into the cortex. (B) scRNA-seq feature plots show *CXCR4*, *CXCL12* (SDF-1) and *ACKR3* (CXCR7) expression in MGEOs at 1 month. (C) Experimental protocol for 2D migration assay in the presence of the CXCR4 antagonist AMD3100 (100 nM). (D) Representative mages of a MGEO from each AMD3100 experimental group after 48 hours post plating (hr pp) at 30 DIV. (E) Quantification of migrated neuron area ± SD in MGEOs 48 hrs pp for 2D migration assay. Eight organoids from 2 differentiations were used per condition and statistical significance was assessed using One-Way ANOVA with repeated measures and Tukey’s multiple comparisons correction. Adjusted P-value for control vs acute AMD3100 treatment conditions was 0.1130 (ns), while chronic condition was statistically significant at 0.0036 (**). (F) Quantification of mDLX-GFP MGEO interneuron area after migration within the CO region of assembloids 48 hours post MEGEO-CO fusion, in the presence of 100 nM AMD3100 or vehicle starting 1 day post fusion. A non-significant trend toward decreased migration is seen. Student T-test statistical comparison between control and AMD3100 treatment organoids (n=4 per group).

**Supplementary Figure S5:**
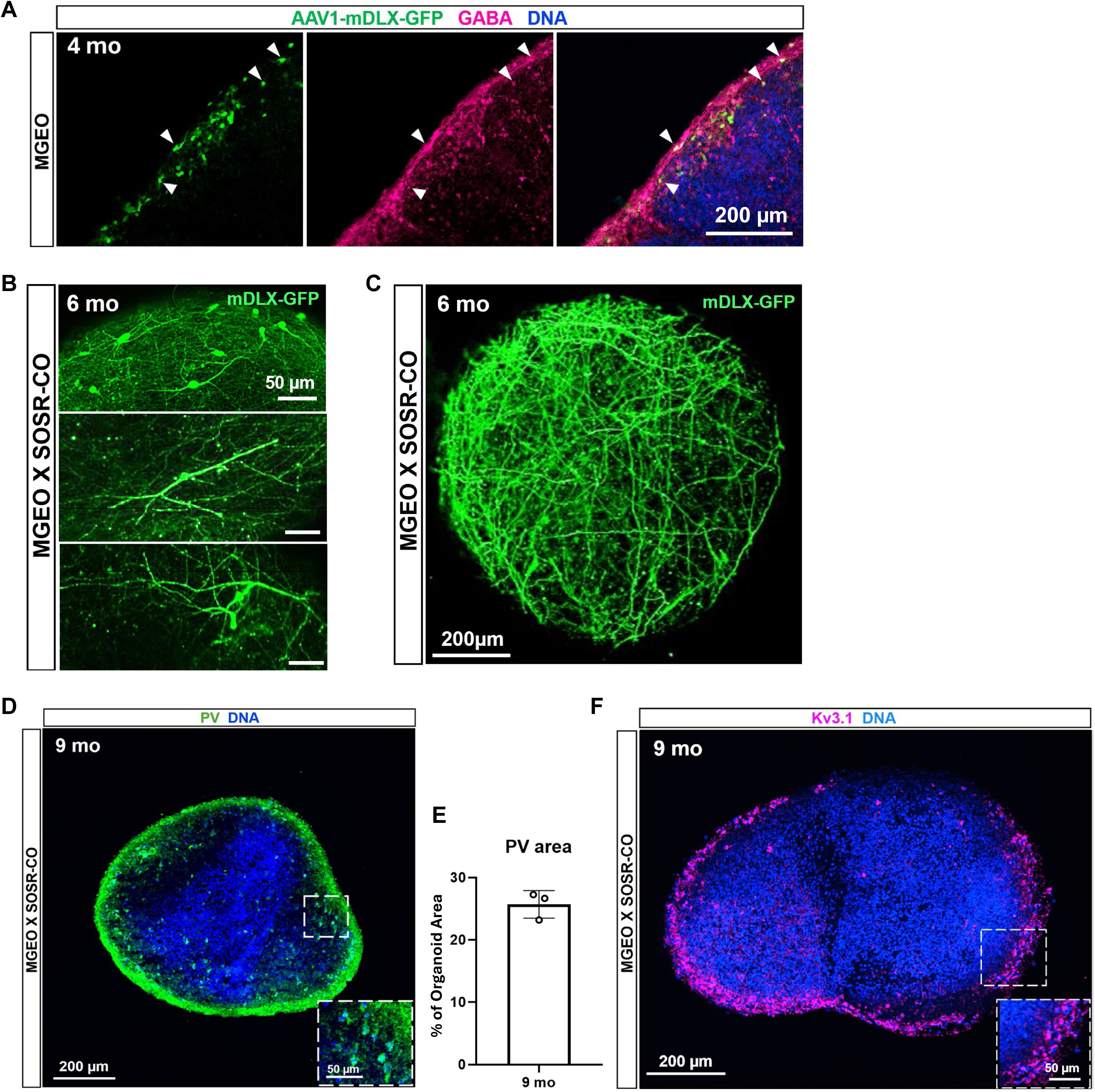
Validation of mDLX-GFP specificity and characterization of interneurons in MGEO-CO assembloids (A) Immunostaining of 4-mo MGEOs shows AAV1-mDLX-GFP virus labeled cells (green) co-expressing GABA (red). Bisbenzimide nuclear stain (DNA) is in blue. (B) mDLX-GFP-labeled MGEO-derived interneurons in assembloids at 6 months (mo) *in vitro* have bipolar and multipolar morphologies typical of MGE-derived interneurons. (C) MGEO-derived interneurons expressing mDLX-GFP on the outer surface of assembloids exhibit extensive process formation by 6 months *in vitro*. (D) PV immunolabeling (green) along the periphery of a 9 month (mo) MGEO-CO assembloid. Bisbenzimide nuclear stain is in blue and inset shows a higher magnification image from the boxed region. (E) Quantification of PV-positive immunolabeling from 9 mo assembloids as percent of organoid area. n=3 assembloids. (F) Immunostaining of 9 mo MGEO-CO assembloid shows expression of the voltage-gated potassium channel K_v_3.1 (fuchsia) along the periphery of the assembloid. Inset shows a higher magnification image from the boxed region.

**Supplementary Figure S6:**
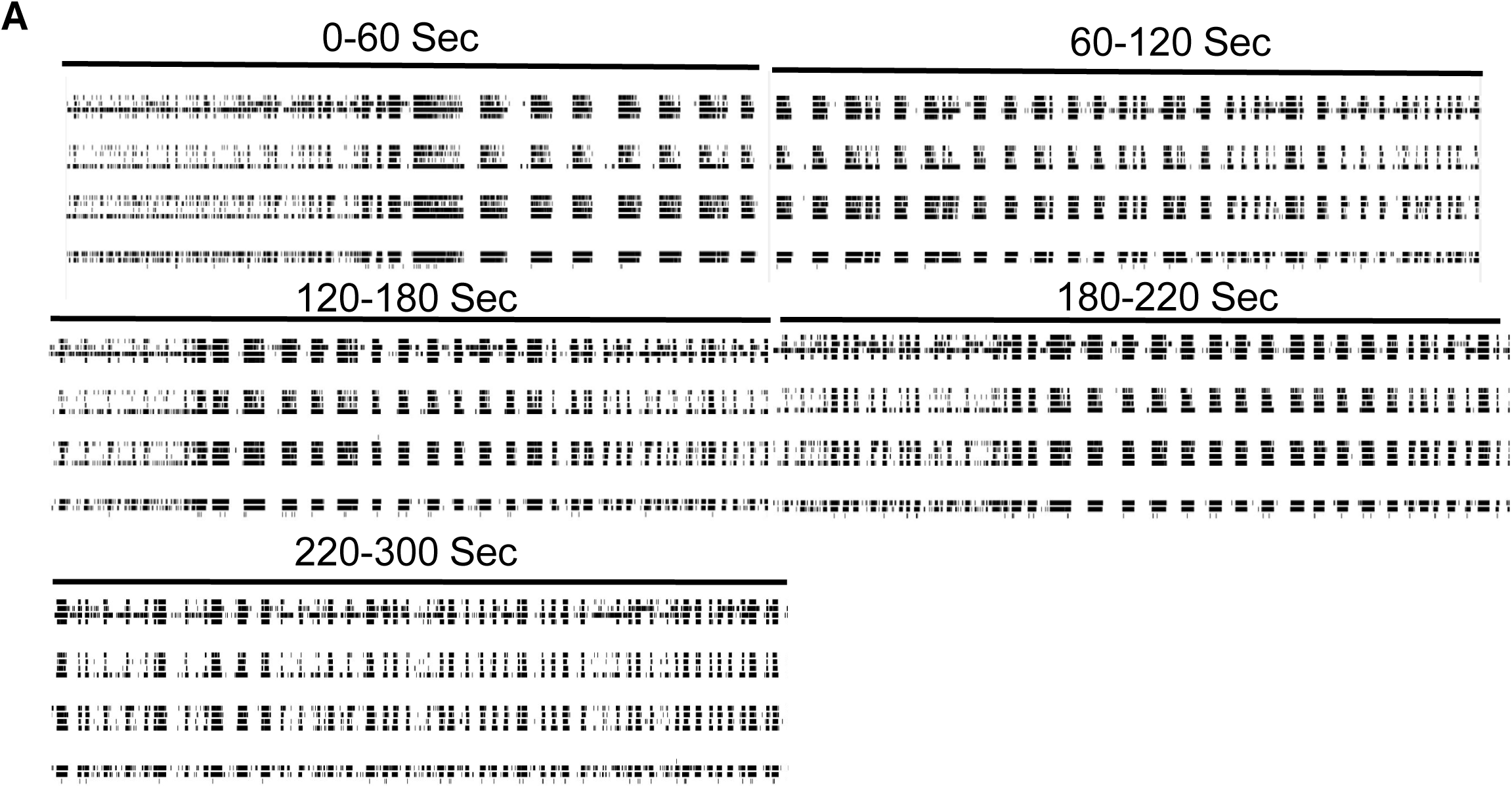
Complexity of MEA network activity (A) Representative continuous 5-minute MEA recording from a 235-day assembloid showing complexity of network bursting activity.

**Supplementary Figure S7:**
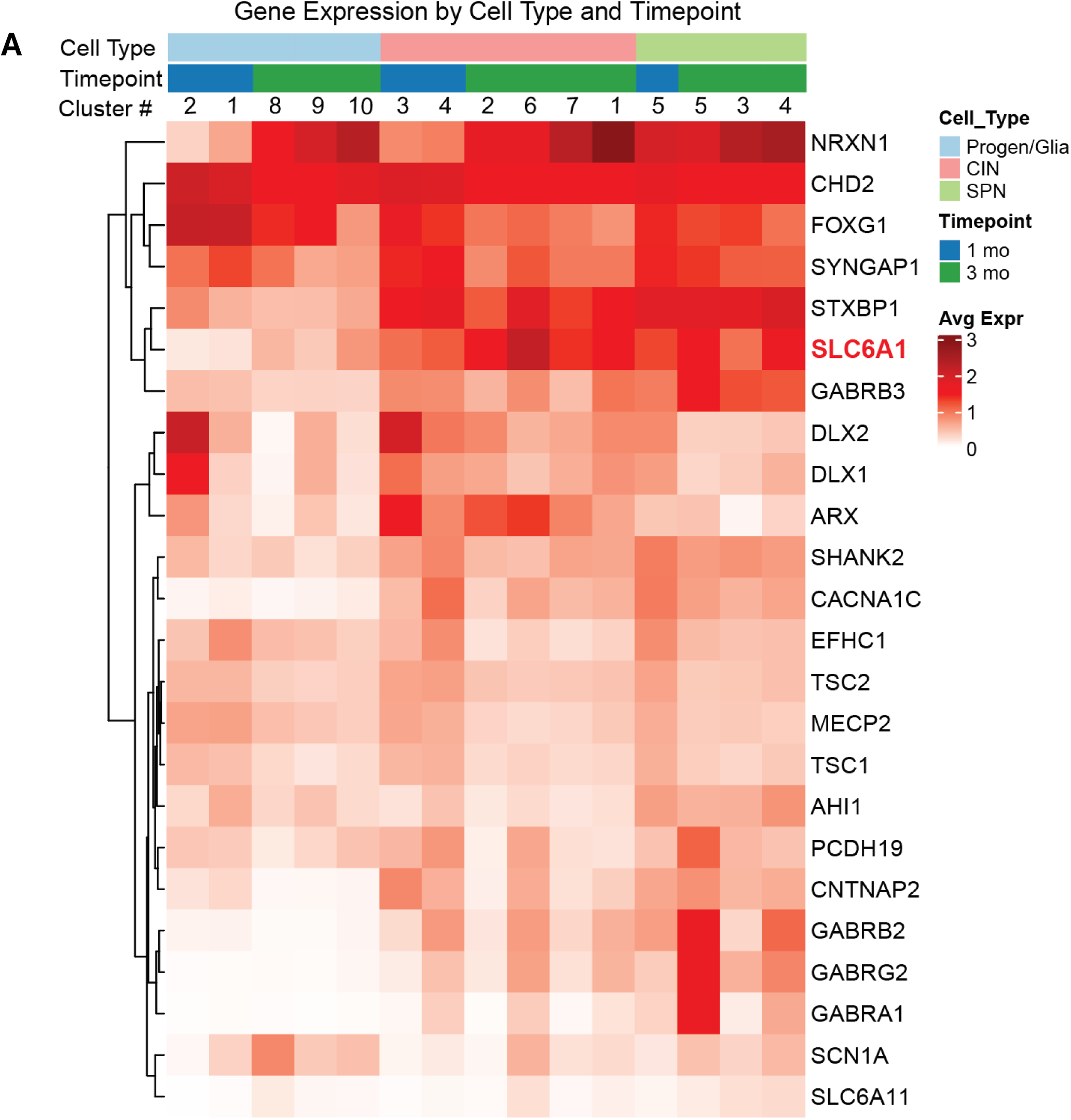
Expression of neurodevelopmental disorder genes in MGEOs (A) Gene expression heat map grouped by cell type, MGEO timepoint, and cluster number showing average expression of a group of genes commonly implicated in various neurodevelopmental disorders. Note high SLC6A1 expression particularly in 3-month interneuron clusters.

**Supplementary Table S1:**
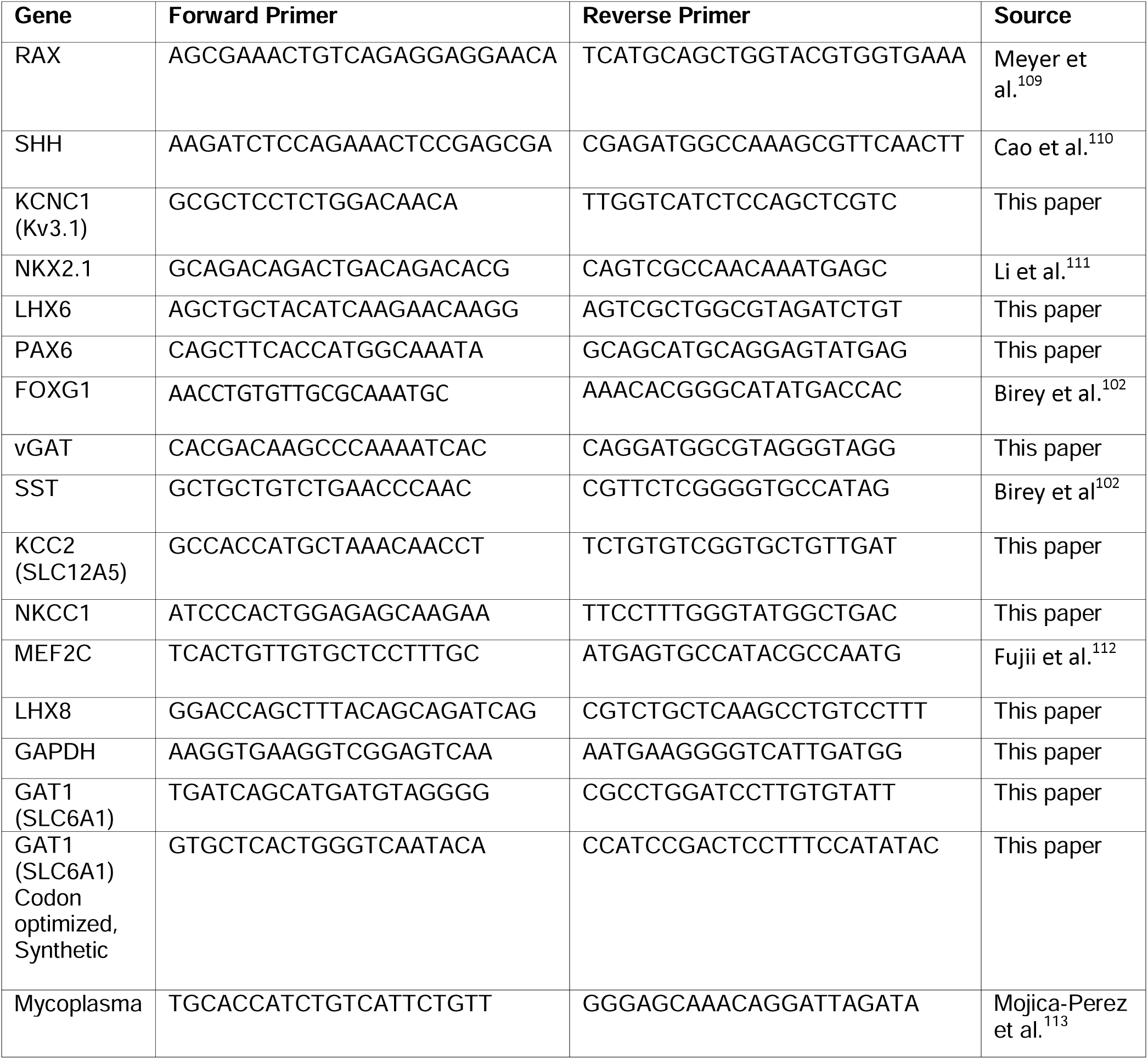
RT-qPCR Primers.

## Notes

### Competing Interest Statement

The authors have declared no competing interest.

### Summary of Updates

New data including a disease model was added to the manuscript. Text was updated to reflect this change. Some main and supplemental figures were adjusted to make room for the new data and data that was inconsequential to the manuscript was removed. One author associated with the removed data was also removed. Title was adjusted to reflect the addition of the disease model.

## References

1. Hájos, N., Pálhalini, J., Mann, E.O., Nèmeth, B., Paulsen, O., and Freund, T.F. (2004). Spike Timing of Distinct Types of GABAergic Interneuron during Hippocampal Gamma Oscillations In Vitro. The Journal of Neuroscience 24, 9127. 10.1523/JNEUROSCI.2113-04.2004.

2. Jacobs, J., Kahana, M.J., Ekstrom, A.D., and Fried, I. (2007). Brain oscillations control timing of single-neuron activity in humans. Journal of Neuroscience 27, 3839–3844. 10.1523/JNEUROSCI.4636-06.2007.

3. Bartos, M., Vida, I., Frotscher, M., Meyer, A., Monyer, H., Geiger, J.R.P., Jonas, P., and Heidelberg (2002). Fast synaptic inhibition promotes synchronized gamma oscillations in hippocampal interneuron networks. Proc Natl Acad Sci U S A 99, 13222–13227. 10.1073/PNAS.192233099.

4. Buzsáki, G., and Chrobak, J.J. (1995). Temporal structure in spatially organized neuronal ensembles: a role for interneuronal networks. Curr Opin Neurobiol 5, 504–510. 10.1016/0959-4388(95)80012-3.

5. Xu, Q., Tam, M., and Anderson, S.A. (2008). Fate mapping Nkx2.1-lineage cells in the mouse telencephalon. Journal of Comparative Neurology 506, 16–29. 10.1002/CNE.21529.

6. Flames, N., Pla, R., Gelman, D.M., Rubenstein, J.L.R., Puelles, L., and Marín, O. (2007). Delineation of multiple subpallial progenitor domains by the combinatorial expression of transcriptional codes. Soc NeuroscienceN Flames, R Pla, DM Gelman, JLR Rubenstein, L Puelles, O MarínJournal of Neuroscience, 2007•Soc Neuroscience. 10.1523/JNEUROSCI.2750-07.2007.

7. Anderson, S.A., Marín, O., Horn, C., Jennings, K., and Rubenstein, J.L.R. (2001). Distinct cortical migrations from the medial and lateral ganglionic eminences. Development 128, 353–363. 10.1242/DEV.128.3.353.

8. Anderson, S.A., Kaznowski, C.E., Horn, C., Rubenstein, J.L.R., and McConnell, S.K. (2002). Distinct origins of neocortical projection neurons and interneurons in vivo. Cereb Cortex 12, 702–709. 10.1093/CERCOR/12.7.702.

9. Wonders, C.P., and Anderson, S.A. (2006). The origin and specification of cortical interneurons. Nature Reviews Neuroscience 2006 7:9 *7*, 687–696. 10.1038/nrn1954.

10. Xu, Q., Cobos, I., De La Cruz, E.D., Rubenstein, J.L., and Anderson, S.A. (2004). Origins of Cortical Interneuron Subtypes. The Journal of Neuroscience 24, 2612. 10.1523/JNEUROSCI.5667-03.2004.

11. Shi, Y., Wang, M., Mi, D., Lu, T., Wang, B., Dong, H., Zhong, S., Chen, Y., Sun, L., Zhou, X., et al. (2021). Mouse and human share conserved transcriptional programs for interneuron development. Science (1979) 374. 10.1126/SCIENCE.ABJ6641.

12. Nicholas, C.R., Chen, J., Tang, Y., Southwell, D.G., Chalmers, N., Vogt, D., Arnold, C.M., Chen, Y.J.J., Stanley, E.G., Elefanty, A.G., et al. (2013). Functional maturation of hPSC-derived forebrain interneurons requires an extended timeline and mimics human neural development. Cell Stem Cell 12, 573–586. 10.1016/j.stem.2013.04.005.

13. Maroof, A.M., Keros, S., Tyson, J.A., Ying, S.W., Ganat, Y.M., Merkle, F.T., Liu, B., Goulburn, A., Stanley, E.G., Elefanty, A.G., et al. (2013). Directed differentiation and functional maturation of cortical interneurons from human embryonic stem cells. Cell Stem Cell 12, 559–572. 10.1016/j.stem.2013.04.008.

14. Keefe, F., Monzón-Sandoval, J., Rosser, A.E., Webber, C., and Li, M. (2023). Single-Cell Transcriptomics Reveals Conserved Regulatory Networks in Human and Mouse Interneuron Development. Int J Mol Sci 24. 10.3390/IJMS24098122.

15. Zhao, Z., Zhang, D., Yang, F., Xu, M., Zhao, S., Pan, T., Liu, C., Liu, Y., Wu, Q., Tu, Q., et al. (2022). Evolutionarily conservative and non-conservative regulatory networks during primate interneuron development revealed by single-cell RNA and ATAC sequencing. Cell Research 2022 32:5 *32*, 425–436. 10.1038/s41422-022-00635-9.

16. Zhao, Y., Flandin, P., Long, J.E., Dela Cuesta, M., Westphal, H., and Rubenstein, J.L.R. (2008). Distinct molecular pathways of development of telencephalic interneuron subtypes revealed through analysis of Lhx6 mutants. Journal of Comparative Neurology 510, 79–99. 10.1002/CNE.21772.

17. Fogarty, M., Grist, M., Gelman, D., Marín, O., Pachnis, V., and Kessaris, N. (2007). Spatial genetic patterning of the embryonic neuroepithelium generates GABAergic interneuron diversity in the adult cortex. Soc NeuroscienceM Fogarty, M Grist, D Gelman, O Marín, V Pachnis, N KessarisJournal of Neuroscience, 2007•Soc Neuroscience 27, 10935–10946. 10.1523/JNEUROSCI.1629-07.2007.

18. Liodis, P., Denaxa, M., Grigoriou, M., Akufo-Addo, C., Yanagawa, Y., and Pachnis, V. (2007). Lhx6 activity is required for the normal migration and specification of cortical interneuron subtypes. Soc NeuroscienceP Liodis, M Denaxa, M Grigoriou, C Akufo-Addo, Y Yanagawa, V PachnisJournal of Neuroscience, 2007•Soc Neuroscience 27, 3078–3089. 10.1523/JNEUROSCI.3055-06.2007.

19. Lim, L., Mi, D., Llorca, A., and Marín, O. (2018). Development and Functional Diversification of Cortical Interneurons. Neuron 100, 294–313. 10.1016/J.NEURON.2018.10.009.

20. Sussel, L., Marin, O., Kimura, S., and Rubenstein, J.L.R. (1999). Loss of Nkx2.1 homeobox gene function results in a ventral to dorsal molecular respecification within the basal telencephalon: evidence for a transformation of the pallidum into the striatum. Development 126, 3359–3370. 10.1242/DEV.126.15.3359.

21. Neves, G., Shah, M.M., Liodis, P., Achimastou, A., Denaxa, M., Roalfe, G., Sesay, A., Walker, M.C., and Pachnis, V. (2013). The LIM homeodomain protein Lhx6 regulates maturation of interneurons and network excitability in the mammalian cortex. Cereb Cortex 23, 1811–1823. 10.1093/CERCOR/BHS159.

22. Vogt, D., Hunt, R.F., Mandal, S., Sandberg, M., Silberberg, S.N., Nagasawa, T., Yang, Z., Baraban, S.C., and Rubenstein, J.L.R. (2014). Lhx6 directly regulates Arx and CXCR7 to determine cortical interneuron fate and laminar position. Neuron 82, 350–364. 10.1016/J.NEURON.2014.02.030.

23. Jakovcevski, I., Mayer, N., and Zecevic, N. (2010). Multiple Origins of Human Neocortical Interneurons Are Supported by Distinct Expression of Transcription Factors. Cerebral Cortex (New York, NY) 21, 1771. 10.1093/CERCOR/BHQ245.

24. Flames, N., Pla, R., Gelman, D.M., Rubenstein, J.L.R., Puelles, L., and Marín, O. (2007). Delineation of Multiple Subpallial Progenitor Domains by the Combinatorial Expression of Transcriptional Codes. Journal of Neuroscience 27, 9682–9695. 10.1523/JNEUROSCI.2750-07.2007.

25. Zhao, Y., Marín, O., Hermesz, E., Powell, A., Flames, N., Palkovits, M., Rubenstein, J.L.R., and Westphal, H. (2003). The LIM-homeobox gene Lhx8 is required for the development of many cholinergic neurons in the mouse forebrain. Proc Natl Acad Sci U S A 100, 9005–9010. 10.1073/PNAS.1537759100.

26. Wichterle, H., Garcia-Verdugo, J.M., Herrera, D.G., and Alvarez-Buylla, A. (1999). Young neurons from medial ganglionic eminence disperse in adult and embryonic brain. Nature Neuroscience 1999 2:5 2, 461–466. 10.1038/8131.

27. Alzu’bi, A., Lindsay, S., Kerwin, J., Looi, S.J., Khalil, F., and Clowry, G.J. (2017). Distinct cortical and sub-cortical neurogenic domains for GABAergic interneuron precursor transcription factors NKX2.1, OLIG2 and COUP-TFII in early fetal human telencephalon. Brain Struct Funct 222, 2309–2328. 10.1007/S00429-016-1343-5.

28. Yu, Y., Zeng, Z., Xie, D., Chen, R., Sha, Y., Huang, S., Cai, W., Chen, W., Li, W., Ke, R., et al. (2021). Interneuron origin and molecular diversity in the human fetal brain. Nat Neurosci 24, 1745–1756. 10.1038/S41593-021-00940-3.

29. Bershteyn, M., Bröer, S., Parekh, M., Maury, Y., Havlicek, S., Kriks, S., Fuentealba, L., Lee, S., Zhou, R., Subramanyam, G., et al. (2023). Human pallial MGE-type GABAergic interneuron cell therapy for chronic focal epilepsy. Cell Stem Cell 30, 1331–1350.e11. 10.1016/J.STEM.2023.08.013.

30. Sloan, S.A., Andersen, J., Pa ca, A.M., Birey, F., and Pa ca, S.P. (2018). Generation and assembly of human brain region–specific three-dimensional cultures. Nature Protocols 2018 13:9 13, 2062–2085. 10.1038/s41596-018-0032-7.

31. Bagley, J.A., Reumann, D., Bian, S., Lévi-Strauss, J., and Knoblich, J.A. (2017). Fused cerebral organoids model interactions between brain regions. Nature Methods 2017 14:7 14, 743–751. 10.1038/nmeth.4304.

32. Birey, F., Andersen, J., Makinson, C.D., Islam, S., Wei, W., Huber, N., Fan, H.C., Metzler, K.R.C., Panagiotakos, G., Thom, N., et al. (2017). Assembly of functionally integrated human forebrain spheroids. Nature 545, 54–59. 10.1038/nature22330.

33. Xiang, Y., Tanaka, Y., Patterson, B., Kang, Y.J., Govindaiah, G., Roselaar, N., Cakir, B., Kim, K.Y., Lombroso, A.P., Hwang, S.M., et al. (2017). Fusion of Regionally Specified hPSC-Derived Organoids Models Human Brain Development and Interneuron Migration. Cell Stem Cell 21, 383–398.e7. 10.1016/j.stem.2017.07.007.

34. Sohal, V.S., Zhang, F., Yizhar, O., and Deisseroth, K. (2009). Parvalbumin neurons and gamma rhythms enhance cortical circuit performance. Nature 459, 698–702. 10.1038/NATURE07991;KWRD=SCIENCE.

35. Jiang, X., Lachance, M., and Rossignol, E. (2016). Involvement of cortical fast-spiking parvalbumin-positive basket cells in epilepsy. Prog Brain Res 226, 81–126. 10.1016/BS.PBR.2016.04.012.

36. Gonzalez-Burgos, G., and Lewis, D.A. (2012). NMDA Receptor Hypofunction, Parvalbumin-Positive Neurons, and Cortical Gamma Oscillations in Schizophrenia. Schizophr Bull 38, 950. 10.1093/SCHBUL/SBS010.

37. Juarez, P., and Martínez Cerdeño, V. (2022). Parvalbumin and parvalbumin chandelier interneurons in autism and other psychiatric disorders. Front Psychiatry 13. 10.3389/FPSYT.2022.913550/FULL.

38. Tidball, A.M., Niu, W., Ma, Q., Takla, T.N., Walker, J.C., Margolis, J.L., Mojica-Perez, S.P., Sudyk, R., Deng, L., Moore, S.J., et al. (2023). Deriving early single-rosette brain organoids from human pluripotent stem cells. Stem Cell Reports 18, 2498–2514. 10.1016/J.STEMCR.2023.10.020.

39. Niu, W., Deng, L., Mojica-Perez, S.P., Tidball, A.M., Sudyk, R., Stokes, K., and Parent, J.M. (2024). Abnormal cell sorting and altered early neurogenesis in a human cortical organoid model of Protocadherin-19 clustering epilepsy. Front Cell Neurosci 18. 10.3389/FNCEL.2024.1339345/FULL.

40. Takla, T.N., Luo, J., Sudyk, R., Huang, J., Walker, J.C., Vora, N.L., Sexton, J.Z., Parent, J.M., and Tidball, A.M. (2023). A Shared Pathogenic Mechanism for Valproic Acid and SHROOM3 Knockout in a Brain Organoid Model of Neural Tube Defects. Cells 12. 10.3390/CELLS12131697.

41. Mo, Z., and Zecevic, N. (2008). Is Pax6 critical for neurogenesis in the human fetal brain? Cereb Cortex 18, 1455–1465. 10.1093/CERCOR/BHM181.

42. Avilés, E.C., Wilson, N.H., and Stoeckli, E.T. (2013). Sonic hedgehog and Wnt: Antagonists in morphogenesis but collaborators in axon guidance. Front Cell Neurosci. 10.3389/FNCEL.2013.00086/FULL.

43. Nóbrega-Pereira, S., Gelman, D., Bartolini, G., Pla, R., Pierani, A., and Marín, O. (2010). Origin and molecular specification of globus pallidus neurons. Soc NeuroscienceS Nóbrega-Pereira, D Gelman, G Bartolini, R Pla, A Pierani, O MarínJournal of Neuroscience, 2010•Soc Neuroscience 30, 2824–2834. 10.1523/JNEUROSCI.4023-09.2010.

44. Du, T., Xu, Q., Ocbina, P.J., and Anderson, S.A. (2008). NKX2.1 specifies cortical interneuron fate by activating Lhx6. Development 135, 1559–1567. 10.1242/DEV.015123.

45. Fragkouli, A., van Wijk, N.V., Lopes, R., Kessaris, N., and Pachnis, V. (2009). LIM homeodomain transcription factor-dependent specification of bipotential MGE progenitors into cholinergic and GABAergic striatal interneurons. Development 136, 3841–3851. 10.1242/DEV.038083.

46. Flandin, P., Kimura, S., Neuroscience, J.R.-J. of, and 2010, undefined (2010). The progenitor zone of the ventral medial ganglionic eminence requires Nkx2-1 to generate most of the globus pallidus but few neocortical interneurons. Soc Neuroscience Journal of Neuroscience 30, 2812–2823. 10.1523/JNEUROSCI.4228-09.2010.

47. Kessaris, N., Fogarty, M., Iannarelli, P., Grist, M., Wegner, M., and Richardson, W.D. (2006). Competing waves of oligodendrocytes in the forebrain and postnatal elimination of an embryonic lineage. Nat Neurosci 9, 173–179. 10.1038/NN1620.

48. Minocha, S., Valloton, D., Ypsilanti, A.R., Fiumelli, H., Allen, E.A., Yanagawa, Y., Marin, O., Chédotal, A., Hornung, J.P., and Lebrand, C. (2015). Nkx2.1-derived astrocytes and neurons together with Slit2 are indispensable for anterior commissure formation. Nat Commun 6. 10.1038/NCOMMS7887.

49. Xu, Q., Cobos, I., De, E., Cruz, L., Rubenstein, J.L., and Anderson, S.A. (2004). Origins of cortical interneuron subtypes. Soc NeuroscienceQ Xu, I Cobos, E De La Cruz, JL Rubenstein, SA AndersonJournal of Neuroscience, 2004•Soc Neuroscience. 10.1523/JNEUROSCI.5667-03.2004.

50. Butt, S.J.B., Fuccillo, M., Nery, S., Noctor, S., Kriegstein, A., Corbin, J.G., and Fishell, G. (2005). The temporal and spatial origins of cortical interneurons predict their physiological subtype. Neuron 48, 591–604. 10.1016/J.NEURON.2005.09.034.

51. Pai, E.L.L., Chen, J., Darbandi, S.F., Cho, F.S., Chen, J., Lindtner, S., Chu, J.S., Paz, J.T., Vogt, D., Paredes, M.F., et al. (2020). Maf and mafb control mouse pallial interneuron fate and maturation through neuropsychiatric disease gene regulation. Elife 9. 10.7554/ELIFE.54903.

52. Allaway, K.C., Gabitto, M.I., Wapinski, O., Saldi, G., Wang, C.Y., Bandler, R.C., Wu, S.J., Bonneau, R., and Fishell, G. (2021). Genetic and epigenetic coordination of cortical interneuron development. Nature 597, 693–697. 10.1038/S41586-021-03933-1.

53. Kaczmarek, L.K., and Zhang, Y. (2017). Kv3 Channels: Enablers of Rapid Firing, Neurotransmitter Release, and Neuronal Endurance. Physiol Rev 97, 1431–1468. 10.1152/PHYSREV.00002.2017.

54. Härtig, W., Brauer, K., and Brückner, G. (1992). Wisteria floribunda agglutinin-labelled nets surround parvalbumin-containing neurons. Neuroreport 3, 869–872. 10.1097/00001756-199210000-00012.

55. Favuzzi, E., Marques-Smith, A., Deogracias, R., Winterflood, C.M., Sánchez-Aguilera, A., Mantoan, L., Maeso, P., Fernandes, C., Ewers, H., and Rico, B. (2017). Activity-Dependent Gating of Parvalbumin Interneuron Function by the Perineuronal Net Protein Brevican. Neuron 95, 639–655.e10. 10.1016/J.NEURON.2017.06.028.

56. Furlanis, E., Dai, M., Leyva Garcia, B., Tran, T., Vergara, J., Pereira, A., Gorissen, B.L., Wills, S., Vlachos, A., Hairston, A., et al. (2025). An enhancer-AAV toolbox to target and manipulate distinct interneuron subtypes. Neuron 113, 1525–1547.e15. 10.1016/J.NEURON.2025.05.002.

57. Birey, F., Andersen, J., Makinson, C.D., Islam, S., Wei, W., Huber, N., Fan, H.C., Metzler, K.R.C., Panagiotakos, G., Thom, N., et al. (2017). Assembly of functionally integrated human forebrain spheroids. Nature 545, 54–59. 10.1038/NATURE22330;TECHMETA=100,13,14,38;SUBJMETA=2182,2571,378,532,631;KWRD=DEVELOPMENT+OF+THE+NERVOUS+SYSTEM,NEURAL+STEM+CELLS.

58. Xiang, Y., Tanaka, Y., Patterson, B., Kang, Y.J., Govindaiah, G., Roselaar, N., Cakir, B., Kim, K.Y., Lombroso, A.P., Hwang, S.M., et al. (2017). Fusion of Regionally Specified hPSC-Derived Organoids Models Human Brain Development and Interneuron Migration. Cell Stem Cell 21, 383–398.e7. 10.1016/j.stem.2017.07.007.

59. Wu, S.J., Sevier, E., Dwivedi, D., Saldi, G.A., Hairston, A., Yu, S., Abbott, L., Choi, D.H., Sherer, M., Qiu, Y., et al. (2023). Cortical somatostatin interneuron subtypes form cell-type-specific circuits. Neuron 111, 2675–2692.e9. 10.1016/J.NEURON.2023.05.032/ASSET/6CFA747C-FE1C-4984-9320-AED19E7E55C3/MAIN.ASSETS/GR6.JPG.

60. Munguba, H., Nikouei, K., Hochgerner, H., Oberst, P., Kouznetsova, A., Ryge, J., Muñoz-Manchado, A.B., Close, J., Batista-Brito, R., Linnarsson, S., et al. (2023). Transcriptional maintenance of cortical somatostatin interneuron subtype identity during migration. Neuron 111, 3590–3603.e5. 10.1016/J.NEURON.2023.07.018.

61. Zhang, C., Xie, Z., and Wang, N. (2025). Single-cell RNA sequencing of adult primate neocortex reveals the regulatory dynamics of neural plasticity. Am J Transl Res 17, 2562. 10.62347/ZEOR5569.

62. Valiente, M., migration, F.M.-C. adhesion &, and 2009, undefined (2009). Migration of cortical interneurons relies on branched leading process dynamics. Taylor & FrancisM Valiente, FJ MartiniCell adhesion & migration, 2009•Taylor & Francis 3, 278–280. 10.4161/cam.3.3.8832.

63. Nadarajah, B., Alifragis, P., Wong, R.O.L., and Parnavelas, J.G. (2003). Neuronal migration in the developing cerebral cortex: Observations based on real-time imaging. Cerebral Cortex 13, 607–611. 10.1093/CERCOR/13.6.607.

64. Pla, R., Borrell, V., Flames, N., Neuroscience, O.M.-J. of, and 2006, undefined (2006). Layer acquisition by cortical GABAergic interneurons is independent of Reelin signaling. Soc NeuroscienceR Pla, V Borrell, N Flames, O MarínJournal of Neuroscience, 2006•Soc Neuroscience. 10.1523/JNEUROSCI.0245-06.2006.

65. Toudji, I., Toumi, A., Chamberland, É., and Rossignol, E. (2023). Interneuron odyssey: molecular mechanisms of tangential migration. Front Neural Circuits 17. 10.3389/FNCIR.2023.1256455/FULL.

66. Stumm, R.K., Zhou, C., Ara, T., Lazarini, F., Dubois-Dalcq, M., Nagasawa, T., Höllt, V., and Schulz, S. (2003). CXCR4 regulates interneuron migration in the developing neocortex. Journal of Neuroscience 23, 5123–5130. 10.1523/JNEUROSCI.23-12-05123.2003.

67. Wang, Y., Li, G., Stanco, A., Long, J.E., Crawford, D., Potter, G.B., Pleasure, S.J., Behrens, T., and Rubenstein, J.L.R. (2011). CXCR4 and CXCR7 Have Distinct Functions in Regulating Interneuron Migration. Neuron 69, 61–76. 10.1016/j.neuron.2010.12.005.

68. Dimidschstein, J., Chen, Q., Tremblay, R., Rogers, S.L., Saldi, G.A., Guo, L., Xu, Q., Liu, R., Lu, C., Chu, J., et al. (2016). A viral strategy for targeting and manipulating interneurons across vertebrate species. Nat Neurosci 19, 1743–1749. 10.1038/NN.4430.

69. Trujillo, C.A., Gao, R., Negraes, P.D., Gu, J., Buchanan, J., Preissl, S., Wang, A., Wu, W., Haddad, G.G., Chaim, I.A., et al. (2019). Complex oscillatory waves emerging from cortical organoids model early human brain network development. Cell Stem Cell 25, 558. 10.1016/J.STEM.2019.08.002.

70. Fattorini, G., Melone, M., and Conti, F. (2020). A Reappraisal of GAT-1 Localization in Neocortex. Front Cell Neurosci 14, 1–6. 10.3389/fncel.2020.00009.

71. Conti, F., Minelli, A., and Melone, M. (2004). GABA transporters in the mammalian cerebral cortex: Localization, development and pathological implications. Brain Res Rev 45, 196–212. 10.1016/j.brainresrev.2004.03.003.

72. Minelli, A., Brecha, N.C., Karschin, C., DeBiasi, S., and Conti, F. (1995). GAT-1, a high-affinity GABA plasma membrane transporter, is localized to neurons and astroglia in the cerebral cortex. J Neurosci 15, 7734–7746. 10.1523/JNEUROSCI.15-11-07734.1995.

73. Carvill, G.L., McMahon, J.M., Schneider, A., Zemel, M., Myers, C.T., Saykally, J., Nguyen, J., Robbiano, A., Zara, F., Specchio, N., et al. (2015). Mutations in the GABA transporter SLC6A1 cause epilepsy with myoclonic-atonic seizures. Am J Hum Genet 96, 808–815. 10.1016/j.ajhg.2015.02.016.

74. Cai, K., Wang, J., Eissman, J., Wang, J., Nwosu, G., Shen, W., Liang, H.C., Li, X.J., Zhu, H.X., Yi, Y.H., et al. (2019). A missense mutation in SLC6A1 associated with Lennox-Gastaut syndrome impairs GABA transporter 1 protein trafficking and function. Exp Neurol 320, 112973. 10.1016/j.expneurol.2019.112973.

75. Mattison, K.A., Butler, K.M., Inglis, G.A.S., Dayan, O., Boussidan, H., Bhambhani, V., Philbrook, B., da Silva, C., Alexander, J.J., Kanner, B.I., et al. (2018). SLC6A1 variants identified in epilepsy patients reduce γ-aminobutyric acid transport. Epilepsia 59, e135–e141. 10.1111/epi.14531.

76. Satterstrom, F.K., Walters, R.K., Singh, T., Wigdor, E.M., Lescai, F., Demontis, D., Kosmicki, J.A., Grove, J., Stevens, C., Bybjerg-Grauholm, J., et al. (2019). Autism spectrum disorder and attention deficit hyperactivity disorder have a similar burden of rare protein-truncating variants. Nature Neuroscience 2019 22:12 22, 1961–1965. 10.1038/s41593-019-0527-8.

77. Tidball, A.M., Dang, L.T., Glenn, T.W., Kilbane, E.G., Klarr, D.J., Margolis, J.L., Uhler, M.D., and Parent, J.M. (2017). Rapid Generation of Human Genetic Loss-of-Function iPSC Lines by Simultaneous Reprogramming and Gene Editing. Stem Cell Reports 9, 725–731. 10.1016/J.STEMCR.2017.07.003.

78. Cuzon, V.C., Yeh, P.W., Cheng, Q., and Yeh, H.H. (2006). Ambient GABA promotes cortical entry of tangentially migrating cells derived from the medial ganglionic eminence. Cereb Cortex 16, 1377–1388. 10.1093/CERCOR/BHJ084.

79. Inada, H., Watanabe, M., Uchida, T., Ishibashi, H., Wake, H., Nemoto, T., Yanagawa, Y., Fukuda, A., and Nabekura, J. (2011). GABA Regulates the Multidirectional Tangential Migration of GABAergic Interneurons in Living Neonatal Mice. PLoS One 6, e27048. 10.1371/JOURNAL.PONE.0027048.

80. Kilb, W., Kirischuk, S., and Luhmann, H.J. (2013). Role of tonic GABAergic currents during pre-and early postnatal rodent development. Front Neural Circuits 7, 59343. 10.3389/FNCIR.2013.00139/FULL.

81. Trujillo, C.A., Gao, R., Negraes, P.D., Gu, J., Buchanan, J., Preissl, S., Wang, A., Wu, W., Haddad, G.G., Chaim, I.A., et al. (2019). Complex Oscillatory Waves Emerging from Cortical Organoids Model Early Human Brain Network Development. Cell Stem Cell 25, 558–569.e7. 10.1016/j.stem.2019.08.002.

82. Melone, M., Ciappelloni, S., and Conti, F. (2014). Plasma membrane transporters GAT-1 and GAT-3 contribute to heterogeneity of GABAergic synapses in neocortex. Front Neuroanat 8. 10.3389/FNANA.2014.00072.

83. Inda, M.C., DeFelipe, J., and Muñoz, A. (2007). The distribution of chandelier cell axon terminals that express the GABA plasma membrane transporter GAT-1 in the human neocortex. Cereb Cortex 17, 2060–2071. 10.1093/CERCOR/BHL114.

84. Wang, J., Poliquin, S., Mermer, F., Eissman, J., Delpire, E., Wang, J., Shen, W., Cai, K., Li, B.M., Li, Z.Y., et al. (2020). Endoplasmic reticulum retention and degradation of a mutation in SLC6A1 associated with epilepsy and autism. Mol Brain 13, 1–15. 10.1186/s13041-020-00612-6.

85. Mermer, F., Poliquin, S., Rigsby, K., Rastogi, A., Shen, W., Romero-Morales, A., Nwosu, G., McGrath, P., Demerast, S., Aoto, J., et al. (2021). Common molecular mechanisms of SLC6A1 variant-mediated neurodevelopmental disorders in astrocytes and neurons. Brain 144, 2499. 10.1093/BRAIN/AWAB207.

86. Piniella, D., Canseco, A., Vidal, S., Xiol, C., Díaz de Bustamante, A., Martí-Carrera, I., Armstrong, J., Bastolla, U., and Zafra, F. (2023). Experimental and Bioinformatic Insights into the Effects of Epileptogenic Variants on the Function and Trafficking of the GABA Transporter GAT-1. Int J Mol Sci 24, 955. 10.3390/IJMS24020955/S1.

87. Cope, D.W., Di Giovanni, G., Fyson, S.J., Orbán, G., Errington, A.C., Lrincz, M.L., Gould, T.M., Carter, D.A., and Crunelli, V. (2009). Enhanced tonic GABAA inhibition in typical absence epilepsy. Nature Medicine 2009 15:12 15, 1392–1398. 10.1038/nm.2058.

88. Chiu, C.S., Brickley, S., Jensen, K., Southwell, A., Mckinney, S., Cull-Candy, S., Mody, I., and Lester, H.A. (2005). GABA Transporter Deficiency Causes Tremor, Ataxia, Nervousness, and Increased GABA-Induced Tonic Conductance in Cerebellum. Journal of Neuroscience 25, 3234–3245. 10.1523/JNEUROSCI.3364-04.2005.

89. Guo, W., Rioux, M., Shaffo, F., Hu, Y., Yu, Z., Xing, C., and Gray, S.J. (2025). AAV9/SLC6A1 gene therapy rescues abnormal EEG patterns and cognitive behavioral deficiencies in Slc6a1–/– mice. J Clin Invest 135. 10.1172/JCI182235.

90. Qi Hu, R., and Davies, J.A. (1997). Tiagabine hydrochloride, an inhibitor of γ-aminobutyric acid (GABA) uptake, induces cortical depolarizations in vitro. Brain Res 753, 260–268. 10.1016/S0006-8993(97)00013-9.

91. Davies, J.A., and Shakesby, A. (1999). Blockade of GABA uptake potentiates GABA-induced depolarizations in adult mouse cortical slices. Neurosci Lett 266, 201–204. 10.1016/S0304-3940(99)00292-X.

92. Matthews, E., Rahnama-Vaghef, A., and Eskandari, S. (2009). Inhibitors of the γ-Aminobutyric Acid Transporter 1 (GAT1) Do Not Reveal a Channel Mode of Conduction. Neurochem Int 55, 732. 10.1016/J.NEUINT.2009.07.005.

93. Jursky, F., and Nelson, N. (1996). Developmental expression of GABA transporters GAT1 and GAT4 suggests involvement in brain maturation. J Neurochem 67, 857–867. 10.1046/J.1471-4159.1996.67020857.X.

94. Ji, L., Xu, Y., and Tang, N. (2021). Solute carrier family 6 member 1 promoting the proliferation and invasion of breast cancer cells through affecting the phosphorylation of PI3K/Akt signaling pathway. Acta Anatomica Sinica 52, 759–766. 10.16098/J.ISSN.0529-1356.2021.05.013.

95. Zhao, Y., Zhou, X., He, Y., and Liao, C. (2018). SLC6A1-miR133a-CDX2 loop regulates SK-OV-3 ovarian cancer cell proliferation, migration and invasion. Oncol Lett 16, 4977–4983. 10.3892/OL.2018.9273/ABSTRACT.

96. Nascimento, M.A., Biagiotti, S., Herranz-Pérez, V., Santiago, S., Bueno, R., Ye, C.J., Abel, T.J., Zhang, Z., Rubio-Moll, J.S., Kriegstein, A.R., et al. (2023). Protracted neuronal recruitment in the temporal lobes of young children. Nature 2023 626:8001 626, 1056–1065. 10.1038/s41586-023-06981-x.

97. Paredes, M.F., James, D., Gil-Perotin, S., Kim, H., Cotter, J.A., Ng, C., Sandoval, K., Rowitch, D.H., Xu, D., McQuillen, P.S., et al. (2016). Extensive migration of young neurons into the infant human frontal lobe. Science (1979) *354*. 10.1126/SCIENCE.AAF7073;JOURNAL:JOURNAL:SCIENCE;ISSUE:ISSUE:DOI.

98. Sanai, N., Nguyen, T., Ihrie, R.A., Mirzadeh, Z., Tsai, H.H., Wong, M., Gupta, N., Berger, M.S., Huang, E., Garcia-Verdugo, J.M., et al. (2011). Corridors of migrating neurons in the human brain and their decline during infancy. Nature 2011 478:7369 478, 382–386. 10.1038/nature10487.

99. Furlanis, E., Dai, M., Leyva Garcia, B., Tran, T., Vergara, J., Pereira, A., Gorissen, B.L., Wills, S., Vlachos, A., Hairston, A., et al. (2025). An enhancer-AAV toolbox to target and manipulate distinct interneuron subtypes. Neuron 113, 1525–1547.e15. 10.1016/j.neuron.2025.05.002.

100. Dimidschstein, J., Chen, Q., Tremblay, R., Rogers, S.L., Saldi, G.A., Guo, L., Xu, Q., Liu, R., Lu, C., Chu, J., et al. (2016). A viral strategy for targeting and manipulating interneurons across vertebrate species. Nat Neurosci 19, 1743–1749. 10.1038/nn.4430.

101. Xiang, Y., Tanaka, Y., Patterson, B., Kang, Y.J., Govindaiah, G., Roselaar, N., Cakir, B., Kim, K.Y., Lombroso, A.P., Hwang, S.M., et al. (2017). Fusion of Regionally Specified hPSC-Derived Organoids Models Human Brain Development and Interneuron Migration. Cell Stem Cell 21, 383–398.e7. 10.1016/j.stem.2017.07.007.

102. Birey, F., Andersen, J., Makinson, C.D., Islam, S., Wei, W., Huber, N., Fan, H.C., Metzler, K.R.C., Panagiotakos, G., Thom, N., et al. (2017). Assembly of functionally integrated human forebrain spheroids. Nature 545, 54–59. 10.1038/nature22330.

103. Tidball, A.M., Dang, L.T., Glenn, T.W., Kilbane, E.G., Klarr, D.J., Margolis, J.L., Uhler, M.D., and Parent, J.M. (2017). Rapid Generation of Human Genetic Loss-of-Function iPSC Lines by Simultaneous Reprogramming and Gene Editing. Stem Cell Reports 9. 10.1016/j.stemcr.2017.07.003.

104. Tidball, A.M., Niu, W., Ma, Q., Takla, T.N., Walker, J.C., Margolis, J.L., Mojica-Perez, S.P., Sudyk, R., Deng, L., Moore, S.J., et al. (2023). Deriving early single-rosette brain organoids from human pluripotent stem cells. Stem Cell Reports 18, 2498–2514. 10.1016/j.stemcr.2023.10.020.

105. Mojica-Perez, S., Stokes, K., Jacobs, S., Huang, J., Vaid, S., Yuan, Y., Pearson, C.A., Montes, D., Tidball, A., VanHeyningen, D., et al. (2025). Cryopreservation of Human Cortical Organoids Using Vitrification. Biorxiv Preprint. 10.1101/2025.04.08.647634.

106. Thomson, J.A. (1998). Embryonic stem cell lines derived from human blastocysts. Science (1979) *282*. 10.1126/science.282.5391.1145.

107. Schindelin, J., Arganda-Carreras, I., Frise, E., Kaynig, V., Longair, M., Pietzsch, T., Preibisch, S., Rueden, C., Saalfeld, S., Schmid, B., et al. (2012). Fiji: An open-source platform for biological-image analysis. Preprint, 10.1038/nmeth.2019.

108. Hao, Y., Hao, S., Andersen-Nissen, E., Mauck, W.M., Zheng, S., Butler, A., Lee, M.J., Wilk, A.J., Darby, C., Zager, M., et al. (2021). Integrated analysis of multimodal single-cell data. Cell 184, 3573–3587.e29. 10.1016/j.cell.2021.04.048.

109. Meyer, J.S., Shearer, R.L., Capowski, E.E., Wright, L.S., Wallace, K.A., Mcmillan, E.L., Zhang, S.-C., and Gamm, D.M. (2009). Modeling early retinal development with human embryonic and induced pluripotent stem cells. Proc. Natl. Acad. Sci. 106, 16698–16703.

110. Cao, S.Y., Hu, Y., Chen, C., Yuan, F., Xu, M., Li, Q., Fang, K.H., Chen, Y., and Liu, Y. (2017). Enhanced derivation of human pluripotent stem cell-derived cortical glutamatergic neurons by a small molecule. Sci Rep 7. 10.1038/s41598-017-03519-w.

111. Li, Y., Eggermont, K., Vanslembrouck, V., and Verfaillie, C.M. (2013). NKX2-1 activation by SMAD2 signaling after definitive endoderm differentiation in human embryonic stem cell. Stem Cells Dev 22, 1433–1442. 10.1089/scd.2012.0620.

112. Fujii, T., Murata, K., Mun, S.H., Bae, S., Lee, Y.J., Pannellini, T., Kang, K., Oliver, D., Park-Min, K.H., and Ivashkiv, L.B. (2021). MEF2C regulates osteoclastogenesis and pathologic bone resorption via c-FOS. Bone Res 9. 10.1038/s41413-020-00120-2.

113. Mojica-Perez, S., Stokes, K., Jacobs, S., Huang, J., Vaid, S., Yuan, Y., Pearson, C.A., Montes, D., Tidball, A., VanHeyningen, D., et al. (2025). Cryopreservation of Human Cortical Organoids Using Vitrification. bioRxiv, 2025.04.08.647634. 10.1101/2025.04.08.647634.

114. Tidball, A.M., Lopez-Santiago, L.F., Yuan, Y., Glenn, T.W., Margolis, J.L., Clayton Walker, J., Kilbane, E.G., Miller, C.A., Martina Bebin, E., Scott Perry, M., et al. (2020). Variant-specific changes in persistent or resurgent sodium current in SCN8A-related epilepsy patient-derived neurons. Brain 143, 3025–3040. 10.1093/BRAIN/AWAA247.

115. Xue, X., Kim, Y.S., Ponce-Arias, A.I., O’Laughlin, R., Yan, R.Z., Kobayashi, N., Tshuva, R.Y., Tsai, Y.H., Sun, S., Zheng, Y., et al. (2024). A patterned human neural tube model using microfluidic gradients. Nature 628, 391–399. 10.1038/S41586-024-07204-7.

116. Yuan, Y., O’Malley, H.A., Smaldino, M.A., Bouza, A.A., Hull, J.M., and Isom, L.L. (2019). Delayed maturation of GABAergic signaling in the Scn1a and Scn1b mouse models of Dravet Syndrome. Sci Rep 9. 10.1038/S41598-019-42191-0.

117. Yuan, Y., Lopez-Santiago, L., Denomme, N., Chen, C., O’Malley, H.A., Hodges, S.L., Ji, S., Han, Z., Christiansen, A., and Isom, L.L. (2024). Antisense oligonucleotides restore excitability, GABA signalling and sodium current density in a Dravet syndrome model. Brain 147, 1231–1246. 10.1093/BRAIN/AWAD349.

118. Lopez-Santiago, L.F., Yuan, Y., Wagnon, J.L., Hull, J.M., Frasier, C.R., O’Malley, H.A., Meisler, M.H., and Isom, L.L. (2017). Neuronal hyperexcitability in a mouse model of SCN8A epileptic encephalopathy. Proc Natl Acad Sci U S A 114, 2383–2388. 10.1073/PNAS.1616821114.

